# The Halastavi árva virus intergenic region IRES promotes translation by the simplest possible initiation mechanism

**DOI:** 10.1101/2020.10.20.347120

**Authors:** Irina S. Abaeva, Quentin Vicens, Anthony Bochler, Heddy Soufari, Angelita Simonetti, Tatyana V. Pestova, Yaser Hashem, Christopher U. T. Hellen

## Abstract

Dicistrovirus intergenic region internal ribosomal entry sites (IGR IRES) do not require initiator tRNA, an AUG codon or initiation factors, and jumpstart translation from the middle of the elongation cycle via formation of IRES/80S complexes resembling the pre-translocation state. eEF2 then translocates the [codon-anticodon]-mimicking pseudoknot I (PKI) from ribosomal A to P sites, bringing the first sense codon into the decoding center. Halastavi árva virus (HalV) contains an IGR that is related to previously described IGR IRESs, but lacks domain 2, which enables these IRESs to bind to individual 40S ribosomal subunits. By employing *in vitro* reconstitution and cryo-electron microscopy, we now report that the HalV IGR IRES functions by the simplest initiation mechanism that involves binding to 80S ribosomes such that PKI is placed in the P site, so that the A site contains the first codon that is directly accessible for decoding without prior eEF2-mediated translocation of PKI.

## INTRODUCTION

The canonical process of translation initiation in eukaryotes begins with separated ribosomal subunits and involves the coordinated activities of >10 eukaryotic initiation factors (eIFs) (Jackson et al., 2010). In outline, the 43S preinitiation complex, comprising the 40S ribosomal subunit, initiator tRNA and eIFs 2, 3, 1 and 1A is recruited to the capped 5’ end of mRNA by eIF4F, eIF4A and eIF4B and scans downstream to the initiation codon, where it stops and forms a 48S initiation complex with established codon-anticodon base-pairing in the ribosomal P site. After recognition of the initiation codon, eIF5 and eIF5B promote dissociation of factors from the 40S subunit and its joining with a 60S subunit, forming an elongation-competent 80S ribosome. This process is regulated in response to physiological changes. For example, cells respond to viral infection by phosphorylation of eIF2, leading to global down-regulation of translation (Mohr and Sonenberg, 2012).

Although the majority of cellular mRNAs initiate translation by the 5’ end-dependent scanning mechanism, initiation on a substantial proportion of viral mRNAs is mediated by so-called internal ribosomal entry sites (IRESs), which contain cis-elements that mediate 5’ end-independent ribosomal recruitment to internal locations within mRNAs. Viral IRESs are grouped into several major classes, each with its own common structural core, conserved sequence motifs and characteristic factor requirements (Mailliot and Martin, 2017). Although structurally unrelated IRESs use distinct initiation mechanisms, all of them are based on non-canonical interactions of IRESs with canonical components of translational apparatus (e.g. eIF4G, eIF3 or 40S and 60S ribosomal subunits) (e.g. Pestova et al., 1996; Pestova et al., 1998b; Wilson et al., 2000; de Breyne et al., 2009; Imai et al., 2016). As a rule, initiation on viral IRESs requires only a subset of canonical eIFs, which allows them to circumvent various cellular regulatory mechanisms (Jackson et al., 2010; Mailliot and Martin, 2017).

The most streamlined translation mechanism identified to date is used by dicistrovirus intergenic region (IGR) IRESs, which exploit direct interaction with the ribosome to jumpstart translation from the elongation stage, skipping requirements for initiator tRNA, an AUG initiation codon and initiation factors (Sasaki and Nakashima, 2000; Wilson et al., 2000; Jan et al., 2003; Pestova and Hellen, 2003). IGR IRESs are ~190nt-long and form two closely related subclasses, epitomized by Cricket paralysis virus (CrPV) (here, designated type VIa) and Taura syndrome virus (TSV) (type VIb). They fold into three pseudoknots: two nested pseudoknots (PKII and PKIII) and a third pseudoknot (PKI), which forms an independent domain (Kanamori and Nakashima, 2001; Nakashima and Uchiumi, 2008). PKII contains an internal loop (L1.1) that binds to the L1 stalk of the 60S subunit, whereas PKIII has two exposed stem-loops, SLIV and SLV, containing conserved apical motifs that bind to the ribosomal proteins eS25, uS7 and uS11 on the head of the 40S subunit (Muhs et al., 2011; Fernández et al., 2014; Koh et al., 2014). PKI mimics the anticodon stem-loop of tRNA base-paired to a cognate codon (Constantino et al., 2008). The first sense codon is located at the 3’ edge of PKI, making PKI responsible for the correct placement of this codon into the ribosomal decoding center. Type VIa and type VIb IGR IRESs have similar compact structures, but in the latter, PKI contains an additional hairpin (SLIII).

IGR IRESs bind either to individual 40S subunits followed by recruitment of 60S subunits, or directly to 80S ribosomes. In IGR IRES-80S complexes, which alternate between canonical and rotated states, the IRES is located in the intersubunit space, with PKI mimicking the tRNA/mRNA interaction in the decoding center of the ribosomal A site (Fernández et al., 2014; Koh et al., 2014). Thus, before the first sense codon can be decoded, PKI has to be removed from the A site. For this, eEF2-GTP binds to the 80S/IRES complex in the rotated state, and induces translocation of PKI to the P site, thereby bringing the first sense codon into the A site where it is decoded by a cognate eEF1A•GTP/aa-tRNA ternary complex (Yamamoto et al., 2007; Fernández et al., 2014; Muhs et al., 2015; Abeyrathne et al., 2016; Murray et al., 2016). This aa-tRNA is released from eEF1A, accommodates in the A site, and eEF2 then promotes another translocation event, moving aa-tRNA to the P site and the IRES to the E site, thereby yielding a ribosomal complex that is competent to begin elongation. Initiation mediated by IGR IRESs thus proceeds via formation of an IRES/80S complex that mimics the pre-translocation state, and starts in the middle of an elongation cycle.

Advances in metagenomics have revealed a plethora of novel dicistrovirus-like viruses (e.g. Culley et al., 2007; Boros et al., 2011; Shi et al., 2016), some of which contain IGR sequences that are shorter than canonical IGR IRESs and lack SLIV and SLV-like motifs. These differences suggest that if these divergent IGRs function as IRESs, they may initiate translation by novel mechanisms. Here, we identified a distinct class of such IGRs epitomized by the IGR from Halastavi árva virus (HalV), which was isolated from the intestinal contents of freshwater carp (*Cyprinus carpio)* (Boros et al., 2011). HalV has a dicistronic RNA genome with an IRES in the 5’UTR that promotes translation of the ORF1 nonstructural protein precursor (Abaeva et al., 2016) and an IGR preceding ORF2, which encodes the capsid protein precursor. By using biochemical analysis and cryo-electron microscopy (cryo-EM) to characterize the HalV IGR, we determined that it contains an IRES that promotes translation by a novel mechanism that is even simpler than that of previously described IGR IRESs.

## RESULTS

### Factor-independent internal ribosomal entry on the HalV IGR

The homology between HalV and dicistrovirus genomes (Boros et al., 2011) suggests that the HalV IGR also promotes initiation on ORF2 by a non-canonical IRES-dependent mechanism. The stop codon of HalV ORF1 is UAA_6276-6278_. The HalV ORF2 is in frame with the UAA_6382-6384_ stop codon downstream of ORF1, and initiation on ORF2 must therefore start after this triplet. The first AUG triplet in ORF2 is AUG_6397-6399_. To determine the start site and the mechanism of initiation on the HalV ORF2, we employed an *in vitro* reconstitution approach. Ribosomal complexes were assembled from individual mammalian translational components (ribosomal subunits, translation factors and aa-tRNAs) on mRNA comprising a 5’-terminal stable hairpin (ΔG=-25.80 kcal/mol) to block the 5’-end dependent initiation, followed by nt. 6211-7460 of the HalV genome. The ribosomal position was then determined by toe-printing, which involves extension by reverse transcriptase of a primer annealed to the ribosome-bound mRNA. cDNA synthesis is arrested by the leading edge of the 40S subunit yielding characteristic toe-prints +15–17 nt from the P site codon (Figure 1A).

**Figure 1.**
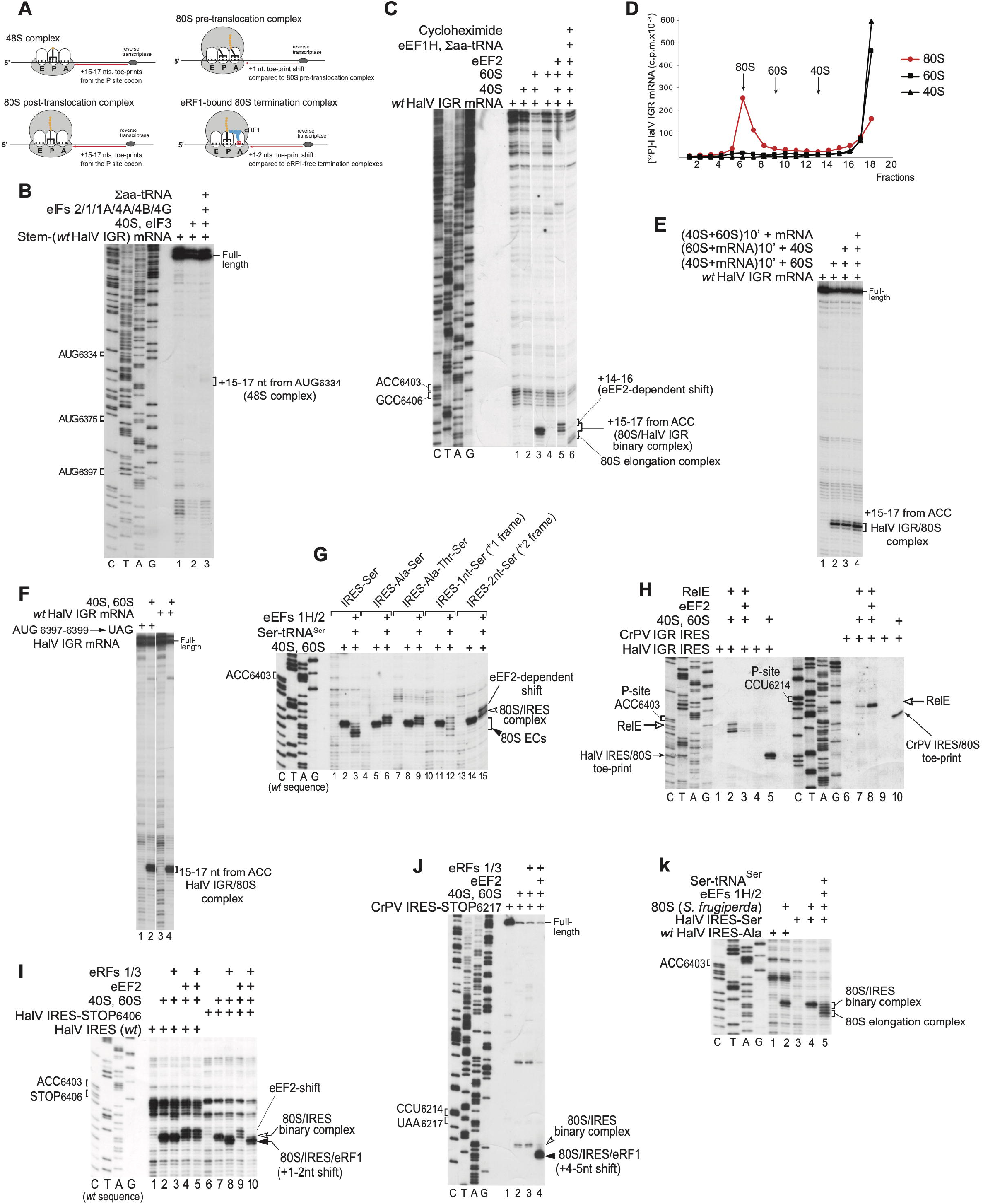
The mechanism of initiation on the HalV IGR IRES. (A) Schematic representation of various ribosomal complexes showing the relative positions of their toeprints. (B) 48S complex formation on HalV IGR mRNA comprising a 5’-terminal stem (ΔG=-25.80 kcal/mol) followed by nt. 6211-7460 of the HalV genome in the presence of 40S subunits, Σaa-tRNA and canonical eIFs, as indicated, assayed by toe-printing. The positions of AUG codons in the HalV IGR and assembled ribosomal complexes are indicated. (C) Direct binding of the HalV IGR mRNA to 80S ribosomes followed by elongation upon addition of eEF1H, eEF2, Σaa-tRNA and cycloheximide, assayed by toe-printing. The positions of the A and P site codons of the HalV IGR IRES are shown on the left. The positions of ribosomal complexes and the eEF2-mediated toe-print shift are indicated on the right. Separation of lanes by white lines indicates that they were juxtaposed from the same gel. (D) Association of [^32^P]-labeled HalV IGR-containing mRNA with individual 40S and 60S ribosomal subunits and 80S ribosomes, assayed by sucrose density gradient centrifugation. (E) Toe-printing analysis of ribosomal association of the HalV IGR IRES depending on the order of incubation of mRNA, 40S and 60S ribosomal subunits. The position of 80S/IRES complexes is indicated. (F) Ribosome-binding activity of the AUG_6397-6399_→UAG stop codon HalV IGR mRNA mutant, assayed by toe-printing. The position of 80S/IRES complexes is indicated. (G) The fidelity of reading frame selection on the HalV IGR IRES investigated by the ability of 80S/IRES complexes formed on GCC_6406-8_ (Ala)→UCU(Ser), ACU_6409-11_(Thr)→UCU(Ser) and AUU_6412-14_(Ile)→UCU(Ser) HalV IGR variants and IGR mutants with UCU(Ser) placed in the +1 or +2 reading frame by insertion of one (G) or two (GC) nts. between ACC6403-5 and UCU_6406-8_(Ser) codons to undergo one-cycle elongation in the presence of eEF1H, eEF2 and Ser-tRNA^Ser^, assayed by toe-printing. The positions of 80S/IRES binary complexes and 80S elongation complexes (80S ECs) are shown on the right. (H) Comparison of the eEF2-dependency of the A site accessibility in 80S ribosomal complexes assembled on HalV and CrPV IGR IRESs, assayed by RelE cleavage. Sites of RelE cleavages were determined by primer-extension. The positions of P site codons, RelE cleavages and 80S/IRES control toe-prints are indicated. (I, J) Comparison of the eEF2-dependency of the A site accessibility in 80S ribosomal complexes formed on HalV and CrPV IGR IRESs by binding of eRF1 and eRF3 to 80S ribosomes assembled on (I) HalV IRES-STOP(UAA) and (J) CrPV IRES-STOP(UAA) mutant mRNAs, assayed by toe-printing. The positions of ribosomal complexes, and P site and Stop codons are indicated. (K) The ability of HalV IGR IRES to form elongation-competent complexes with insect (*Spodoptera frugiperda*) 80S ribosomes, assayed by toe-printing. The positions of the A site codon and ribosomal complexes are indicated. (B, C, G-K) Lanes *C*, *T, A*, and *G* depict CrPV or HalV sequences, as indicated.

In the presence of 40S subunits, Met-tRNA_i_^Met^ and all canonical eIFs, we observed faint toe-prints that could correspond to 48S complexes formed at the out-of-frame AUG_6334_ triplet, but no toe-prints that could be consistent with 48S complex formation at any of ORF2’s in-frame codons (Figure 1B). Strikingly, although in contrast to the CrPV IGR IRES, HalV mRNA did not bind to individual 40S subunits (Figure 1C, lane 2; Figure 1D, black triangles), it bound efficiently to 80S ribosomes, yielding prominent toe-prints ^+^15-^+^17nt downstream of the ACC_6403-6405_ codon in ORF2 (Figure 1C, lane 3; Figure 1D, red circles). To confirm that HalV mRNA binds directly to pre-assembled 80S ribosomes, we performed preincubation experiments. Thus, mRNA was preincubated with 40S or 60S subunits and then 60S or 40S subunits were added, respectively, or 40S and 60S subunits were first preincubated to form 80S ribosomes and then mRNA was added to the mixture. The highest yield of toe-prints ^+^15-^+^17nt downstream of the ACC_6403-6405_ codon was observed in the last case (Figure 1E), indicating that HalV mRNA binds directly to 80S ribosomes. The position of toe-prints suggested that ACC_6403-6405_ and the following ORF2’s GCC_6406-6408_ (Ala) triplet occupied the ribosomal P and A sites, respectively, and that ribosomes did not recognize AUG_6397-6399_, the first AUG in ORF2. Moreover, the AUG_6397-6399_→UAG stop codon mutant HalV mRNA bound 80S ribosomes as efficiently as the *wt* HalV mRNA (Figure 1F). Addition of eEF1H, eEF2 and total aa-tRNA to 80S/HalV mRNA complexes enabled ribosomes to undergo elongation, which was arrested by cycloheximide (Figure 1C, lane 6). Notably, in the absence of eEF1H and aa-tRNA, eEF2 induced a −2nt upward shift of the toe-print (Fig. 1C, lane 5) that may reflect its trapping of the ribosome in the rotated state or with the head in a swiveled position (e.g. Flis et al., 2018). In conclusion, the HalV IGR contains an IRES that binds productively to 80S ribosomes in a factor-independent manner.

To facilitate investigation of the mechanisms of delivery of the first aa-tRNA and initial translocation events on the HalV IGR IRES, the presumed A site GCC_6406-6408_ (Ala) codon was replaced by the UCU_6406-6408_ (Ser) codon (a GCC_6406-6408_(Ala)→UCU(Ser) HalV IGR IRES variant) because *in vitro* transcribed Ser-tRNA^Ser^-AGA is readily available and active in mammalian elongation (Zinoviev et al., 2018). Addition of Ser-tRNA^Ser^, eEF1H and eEF2 to 80S ribosomes bound to a GCC_6406_(Ala)→UCU(Ser) IGR variant resulted in a forward toe-print shift by precisely 3 nucleotides (Figure 1G, lane 3), which would be consistent with one, but not two cycles of translocation as in the case of the CrPV IRES. Such translocation did not occur when the UCU (Ser) codon replaced the second or third triplet downstream of the IRES, and in this case we observed only the eEF2-induced upward toe-print shift (Figure 1G, lanes 6, 9). However, a low level of templated translocation was apparent in the +1 but not in the +2 frame (Figure 1G, lanes 12 and 15), indicating that the fidelity of initiation is high but not absolute. Notably, in the case with the UCU (Ser) codon in the +1 position, we observed not only the +4nt toe-print shift that corresponds to one cycle of translocation, but also a more prominent toe-print at the +2 position indicative of binding of Ser-tRNA^Ser^ (Zinoviev et al., 2015) in the pre-translocation state, reflecting inefficient translocation in this reading frame after delivery of aa-tRNA to the A site.

To confirm that binding of the HalV IGR IRES to 80S ribosomes places the GCC_6406-6408_ triplet directly in the ribosomal A site without a prior eEF2-dependent pseudo-translocation event, we employed the bacterial toxin RelE, which cleaves mRNA in the A-site (Pedersen et al, 2003; Neubauer et al, 2009; Pisareva et al., 2011). RelE efficiently cleaved the 80S-bound HalV IGR IRES within the GCC_6406-6408_ triplet (Figure 1H, lane 2). Importantly, in contrast to cleavage of the CrPV IRES that occurred efficiently only on inclusion of eEF2 (Figure 1H, compare lanes 7 and 8), consistent with the requirement for eEF2 to translocate PKI from the A site (Fernández et al., 2014), cleavage of the HalV IRES not only occurred independently of eEF2 but was even inhibited by it (Figure 1H, lane 3), likely reflecting competition between RelE and eEF2 for binding to the A site. The A-site in HalV IRES-bound 80S ribosomes is therefore vacant and can accept aa-tRNA directly. To further prove that the A-site is vacant, we replaced it by a stop codon. 80S ribosomes associated with the GCC_6406_→stop codon HalV IRES variant efficiently bound eRF1•eRF3 independently of eEF2, evident by characteristic +1-2nt toe-print shift caused by the A site mRNA compaction (Figure 1I, lanes 7 and 8) (Alkalaeva et al., 2006; Brown et al., 2015). In contrast, 80S ribosomes associated with the analogous CrPV IRES mutant did require eEF2 to bind eRF1•eRF3, and binding induced a +4-5nt toe-print shift that included eEF2-dependent initial IRES translocation and mRNA compaction caused by binding of eRF1•eRF3 (Figure 1J; Muhs et al., 2015).

The HalV IGR IRES also efficiently bound to insect (*Spodoptera frugiperda)* 80S ribosomes yielding prominent toe-prints ^+^15-^+^17nt downstream of ACC_6403-6405_ and underwent one cycle of translocation on the GCC_6406-6408_(Ala)→UCU(Ser) HalV IRES variant in the presence of eEFs and Ser-tRNA^Ser^ (Figure 1K).

In conclusion, *in vitro* reconstitution revealed that the HalV IGR contains an IRES which binds directly to 80S ribosomes and jumpstarts translation from the elongation stage by accepting aa-tRNA to the ribosomal A site, containing the Ala codon GCC_6406-6408_. In contrast to dicistrovirus CrPV-like IGR IRESs, this step does not require prior eEF2-mediated translocation of a portion of the IRES from A to P sites. However, similarly to CrPV-like IRESs (e.g. Petrov et al., 2016), the HalV IRES is not wholly specific in determining the reading frame for translation, with a minor fraction of initiation events occurring in the +1 frame.

### Inhibition of ribosomal binding of the HalV IGR IRES by SERPINE1 mRNA binding protein 1 (SERBP1) and eEF2

Although the HalV IGR IRES bound productively to 80S ribosomes assembled *in vitro* from individual salt-washed ribosomal subunits, it did not promote efficient translation in rabbit reticulocyte lysate (RRL) (Figure 2A, lane 5). However, the translational efficiency was increased by addition of assembled *in vitro* vacant 80S ribosomes, and particularly, by preincubation of mRNA with such ribosomes before the addition of both to the translation mixtures (Figure 2A, lanes 6-7). In contrast, translation promoted by the 5’-terminal HalV IRES (Abaeva et al., 2016) was not significantly affected by addition of 80S ribosomes, irrespective of preincubation (Figure 2A, lanes 2-4). Non-programmed 80S ribosomes in RRL, human peripheral blood mononuclear cells and *D. melanogaster* embryonic extracts are associated with SERBP1 and eEF2 (Anger et al., 2013; Zinoviev et al. 2015; Brown et al., 2018; Figure 2B). SERBP1 binds to the head of the 40S subunit and then enters the mRNA-binding channel and follows the mRNA-binding path until the A site where it interacts with domain IV of eEF2 (Anger et al., 2013; Brown et al., 2018). It prompted us to investigate whether ribosomal association with SERBP1 and eEF2 is responsible for the low translational activity of the HalV IGR IRES in RRL. Preincubation of 80S ribosomes with SERBP1 strongly reduced their ability to bind the HalV IGR IRES, whereas preincubation with SERBP1 and eEF2 nearly abrogated formation of 80S/IRES complexes (Figure 2C). Consistently, SERBP1/eEF2-associated native purified RRL 80S ribosomes (Figure 2B) did not bind the HalV IGR IRES (Figure 2D). In contrast, SERBP1 and eEF2 did not affect ribosomal association of the CrPV IRES (Figure 2E). Thus, the presence of SERBP1 and eEF2 on 80S ribosomes specifically prevents their binding to the HalV IGR IRES and is responsible for the low activity of the IRES in RRL.

**Figure 2.**
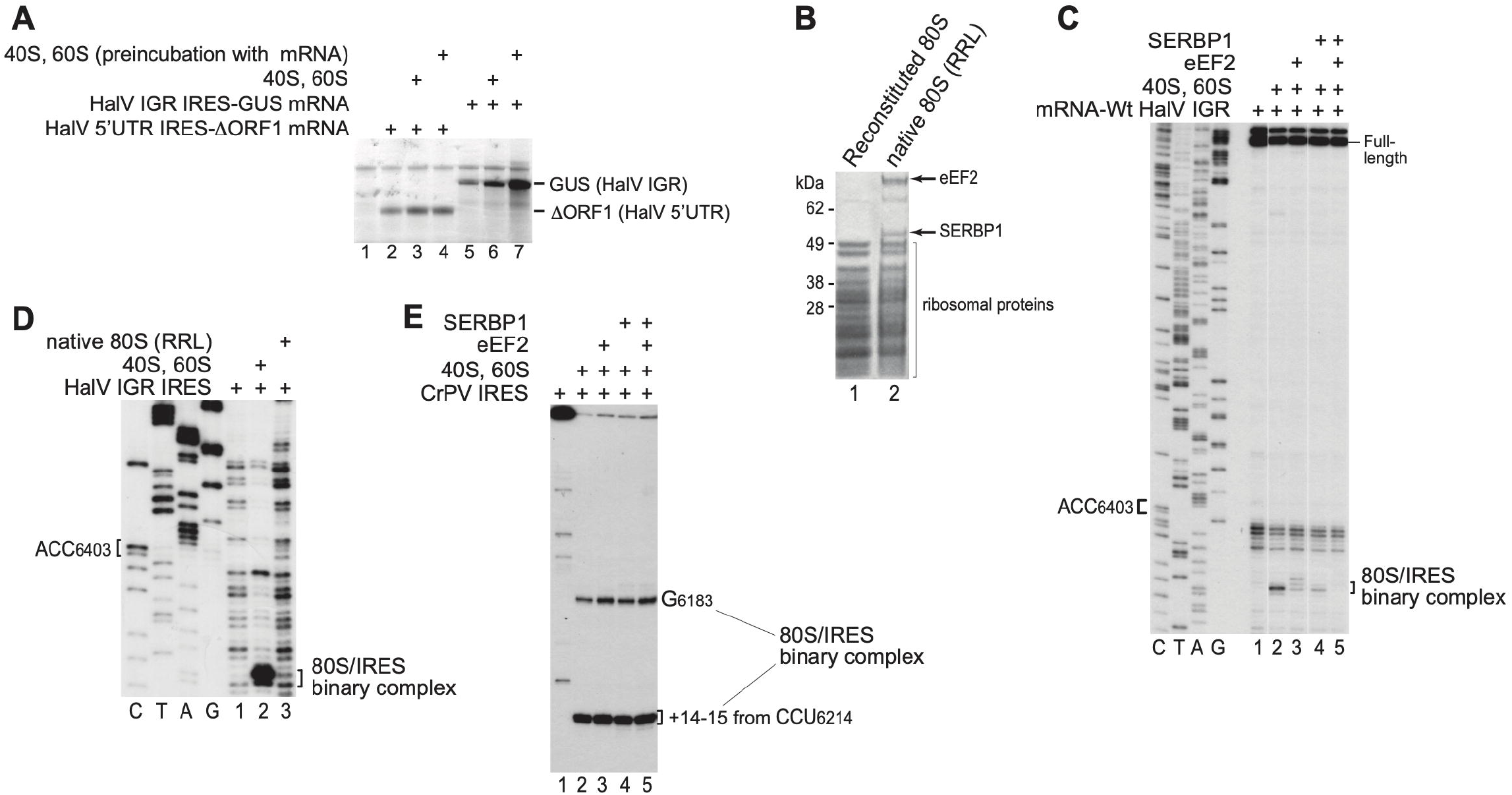
(A) Translation in RRL driven by HalV 5’UTR and IGR IRESs, depending on addition of 40S and 60S subunits with or without their pre-incubation with mRNA. (B) Protein composition of 80S ribosomes reconstituted from individual purified 40S and 60S subunits, and native 80S ribosomes purified from RRL, assayed by SDS-PAGE followed by SYPRO staining. (C) The influence of SERBP1 with/without eEF2 on ribosomal binding of the HalV IGR IRES, assayed by toe-printing. The positions of the A site codon and ribosomal complexes are indicated. Separation of lanes by white lines indicates that they were juxtaposed from the same gel. (D) Comparison of the binding of the HalV IGR IRES to reconstituted and native 80S ribosomes, assayed by toe-printing. The positions of the A site codon and ribosomal complexes are indicated. (C, D) Lanes *C*, *T, A*, and *G* depict HalV sequence. (E) The influence of SERBP1 with/without eEF2 on ribosomal binding of CrPV IGR IRES, assayed by toe-printing. The positions of toe-prints corresponding to 80S/IRES complexes are indicated.

### Structural model of the HalV IGR IRES

To aid understanding of the mechanism of the HalV IGR IRES function, a structural model of the HalV IGR and the adjacent 3’-terminal region of ORF1 (Figure 3A) was derived on the basis of computational analysis (see Materials and Methods) and chemical/enzymatic foot-printing (Figures S1A-B). The 3’-terminal region of ORF1 (nts 6211-6267) forms a large hairpin (Figures 3A and S1C). The HalV IGR consists of two domains that are comparable to domains 1 and 3 of the CrPV-like IGR IRESs (compare Figures 3A and 3B). Domain 1 is represented by a pseudoknot (PKII), comprising helical elements P1.1 (which contains the ORF1 stop codon), P1.2 and P1.3, and internal loops, including loops (e.g. L1.1) with sequences like those in corresponding loops in CrPV-like IGR IRESs (Figures 3A-B and S2). Domain 3 (named in accordance with the nomenclature for CrPV-like IGR IRESs (Costantino and Kieft, 2005)) comprises a second pseudoknot, PKI, at the 3’-border of the IGR and ends with the A-site GCC_6406-6408_ (Ala) codon. There is no equivalent of domain 2 comprising PKIII of CrPV-like IGR IRESs or of the conserved stem-loops IV and V in them that bind to the 40S subunit.

**Figure 3.**
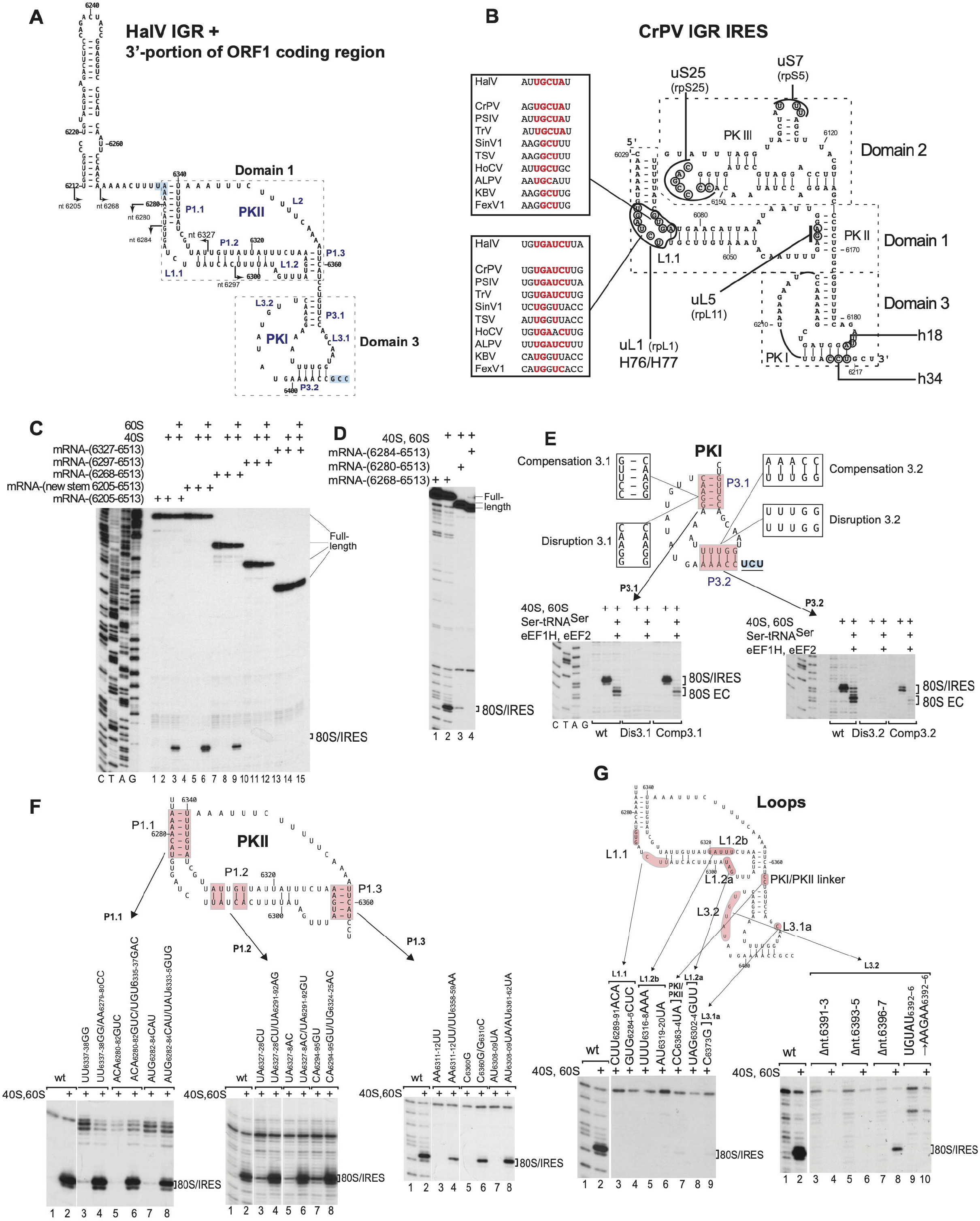
Structure and mutational analysis of the HalV IGR IRES. (A) Model of the HalV IGR IRES and the adjacent 3’-terminal region of ORF1, derived on the basis of computational analysis (see Materials and Methods) and chemical/enzymatic foot-printing (Figures S1A-B). It is annotated to show nucleotides at 20-nt. intervals, IRES domains and secondary structural elements (based on the nomenclature proposed by Constantino and Kieft (2005)), the ORF1 stop codon (UAA_6276_) and the ORF2 A site codon (GCC_6406_) (both boxed blue). Arrows indicate the 5’-borders of truncated HalV IGR mRNAs used in experiments to assay IRES activity in ribosome binding (panels C-D). (B) Model of the CrPV IGR IRES, showing IRES domains and secondary structure elements, nucleotides that interact with ribosomal proteins and elements of 18S and 28S rRNAs, and conserved motifs in the L1.1 loop in dicistrovirus IGR IRESs (Nishiyama et al., 2003; Pfingsten et al., 2006). (C, D) The 5’-terminal border of the HalV IGR IRES assayed by toe-printing of 80S/IRES complex formation. The position of 80S/IRES binary complexes is indicated. Lanes *C, T, A*, and *G* depict HalV sequence. (E, F) Analysis of the influence of disruptive and compensatory substitutions in helical elements of domain 3/PKI (E) and domain 1/PKII (F) on the HalV IGR IRES’s ribosome-binding (E, F) and elongation (E) activity, analyzed by toeprinting. The positions of 80S/IRES binary complexes and 80S ECs are indicated. (G) Analysis of the influence of substitutions in single-stranded elements of the HalV IGR IRES on its ribosome-binding activity, analyzed by toeprinting. The position of 80S/IRES binary complexes is indicated. (F, G) Separation of lanes by white lines indicates that they were juxtaposed from the same gel.

### The HalV IGR IRES epitomizes a novel class of IGR IRESs

Database searches identified HalV-like IGRs in several uncategorized viral genomes from arthropods, including Changjiang picorna-like virus 14 (CPLV14), Shahe arthropod virus 1 (SAV1) (Shi et al., 2016), Kuiper virus and the identical Drellivirus. A partial HalV-like IGR sequence (*P. solanasi* TSA-1) was identified in a Transcriptome shotgun assembly (TSA) sequence from the crustacean *Proasellus solanasi*. The nucleotide sequence of this IRES fragment and the amino acid sequence of the 82 a.a.-long ORF2 fragment are 86% and 57% identical to CPLV14, respectively.

In these genomes, the IGRs are flanked by ORF1 that encodes dicistrovirus-like nonstructural proteins and ORF2 that encodes capsid proteins in the order VP2-VP4-VP3-VP1 that is characteristic of dicistroviruses. Amino acid sequence identity in the 3C protease/3D polymerase segment of ORF1 of these viruses ranges from 32.7 - 71.2% and in ORF2, from 43.2 - 60.2% (Supplementary Tables 1, 2). Identity between ORF1 and ORF2 of these viruses and of representative members of the *Cripavirus* genus (CrPV), *Triatovirus* genus (Triatoma virus) and the two clades of insect- and crustacean-infecting viruses (ABPV and TSV, respectively) in the *Aparavirus* genus of *Dicistroviridae* (www.ictv.global/report/dicistroviridae) did not exceed 27.4% for the 3CD region and 25.1% for the ORF2 region, respectively (Supplementary Tables 1, 2). Phylogenetic analysis of ORF2 amino acid sequences confirmed that HalV, CPLV14, SAV1 and Kuiper virus form a clade, here designated ‘Halárvirus’, that is distinct from the *Aparavirus, Cripavirus* and *Triatovirus* genera (Figure S1D).

The HalV IGR IRES is 129nt. long from the 5’-border of P1.1 to the 3’-border of PKI, and these putative Halávirus IGR IRESs are similar (SAV1: 123nt.; CPLV14: 126nt.; Kuiper virus: 129nt.; Figure S2). By extension of the nomenclature for CrPV-like IGR IRESs (type VIa) and TSV-like IGR IRESs (type VIb), we suggest that HalV-like IGR IRESs be designated type VIc (Figure S2). Pairwise sequence identity between them is high (45-66%) and they all have a HalV IGR-like structure comprising two domains with a large loop in domain 1 instead of a PKIII-like element. Conserved nucleotides are concentrated in the L1 loop, in P3.2 and in the adjacent L3.1 and L3.2 elements of PK1 (Figure S2).

### Mutational analysis of the HalV IRES

Progressive 5’-terminal deletions were made to determine the 5’ border of the HalV IGR IRES (indicated in Figure 3A). Deletion of the ORF1 hairpin and its replacement by a mirror-image version were tolerated, whereas deletion to nt. 6280 and nt. 6284 strongly reduced and abrogated ribosomal binding of the IRES, respectively (Figures 3C-D). The ORF1 hairpin is therefore dispensable, nt. 6268-6405 are sufficient, and the integrity of P1.1 is important for IRES function in ribosome binding. Thus, the HalV IGR IRES is 129nt-long, and is consequently a third smaller than previously characterized IGR IRESs (~190nt).

To verify the HalV IGR IRES model, we performed mutagenesis to confirm the ability of certain regions to form the predicted functionally important secondary structure elements. For this, destabilizing and compensatory mutations were introduced into PKI and PKII. Ribosomal binding and elongation activities of the GCC_6406-6408_(Ala)→UCU(Ser) IRES variant were abrogated by substitutions in P3.1 and P3.2 of PKI, but were restored fully or partially, respectively, by compensatory second-site substitutions (Figure 3E). Similarly, in the case of PKII, IRES function was abrogated by destabilizing substitutions in P1.1 and P1.3 and strongly impaired by destabilizing substitutions in the proximal region of P1.2, but was fully restored by compensatory substitutions (Figure 3F).

Whereas helical elements function to support the structure of the IRES, single-stranded regions might engage in specific interactions with the ribosome to ensure efficient, stable binding or correct orientation of functionally important motifs. Mutational analysis revealed that whereas the long L2 loop (nt 6341-6357) in PKII that replaces PKIII/domain 2 of CrPV-like IGR IRESs was quite tolerant of substitutions (Figure S3), other unpaired elements, like the L1.1, L1.2a and L1.2b loops in PKII, L3.1a and L3.2 loops in PKII, and the PKI/PKII linker had critical functions that were strongly sequence-dependent (Figure 3G). These elements contribute to or are essential for the function of previously described IGR IRESs, and many of them interact with components of the ribosome (Figure 3B). Thus, the L1.1 loop binds to uL1 and H76/H77 of the L1 stalk (e.g. Schüler et al., 2006; Fernández et al., 2014) and is important for ribosomal attachment to and translocation of the IRES (Jang et al., 2009; Pfingsten et al., 2006, 2010), the L3.1a loop interacts with h18 (Schüler et al., 2006) and substitutions in it reduce IRES-mediated translation (Costantino et al., 2008), and the L3.2 loop, which interacts with uS7 (Abeyrathne et al., 206; Acosta-Reyes et al., 2019), plays a role in translocation (Ruehle et al., 2015).

### Structure of the HalV IGR IRES bound to the rabbit 80S ribosome

Cryo-EM of the HalV IRES/rabbit 80S ribosome complex yielded two classes (Figure S4), one with rotated ribosomes and the other with ribosomes in the unrotated (classical) conformation (Figure 4A). Refinement of the rotated class yielded an average resolution of 3.6 Å. However, the 40S subunit is likely to be oscillating between several close rotated conformations, as its densities appear scanter than those of the 60S subunit. The unrotated class appears more sturdy and presents similarly solid cryo-EM densities for both ribosomal subunits, yielding a 3.49 Å resolution reconstruction (Figure 4B). Ribosomes in the unrotated state resembled those in CrPV IGR IRES/80S complexes after the first or second translocation steps, when the IRES is either in the P- or E- site, or the hybrid P/E state (Figure S5) (Muhs et al., 2015; Pisareva et al., 2018). Hence, the head of the 40S subunit is not swiveled as in the complex with the CrPV IRES bound to the A site of the 80S ribosome (Fernández et al., 2014) and in ribosomes in the POST state (Budkevich et al., 2014). The rotated state is similar to that observed in the presence of A-site bound CrPV IRES (Fernández et al., 2014) and is closely related to the PRE-1 state (Figure S5) (Budkevich et al., 2014). In both classes, PKI mimics the anticodon stem-loop of tRNA base-paired to a mRNA codon in the P site, leaving the A site vacant, consistent with the biochemical data described above.

**Figure 4.**
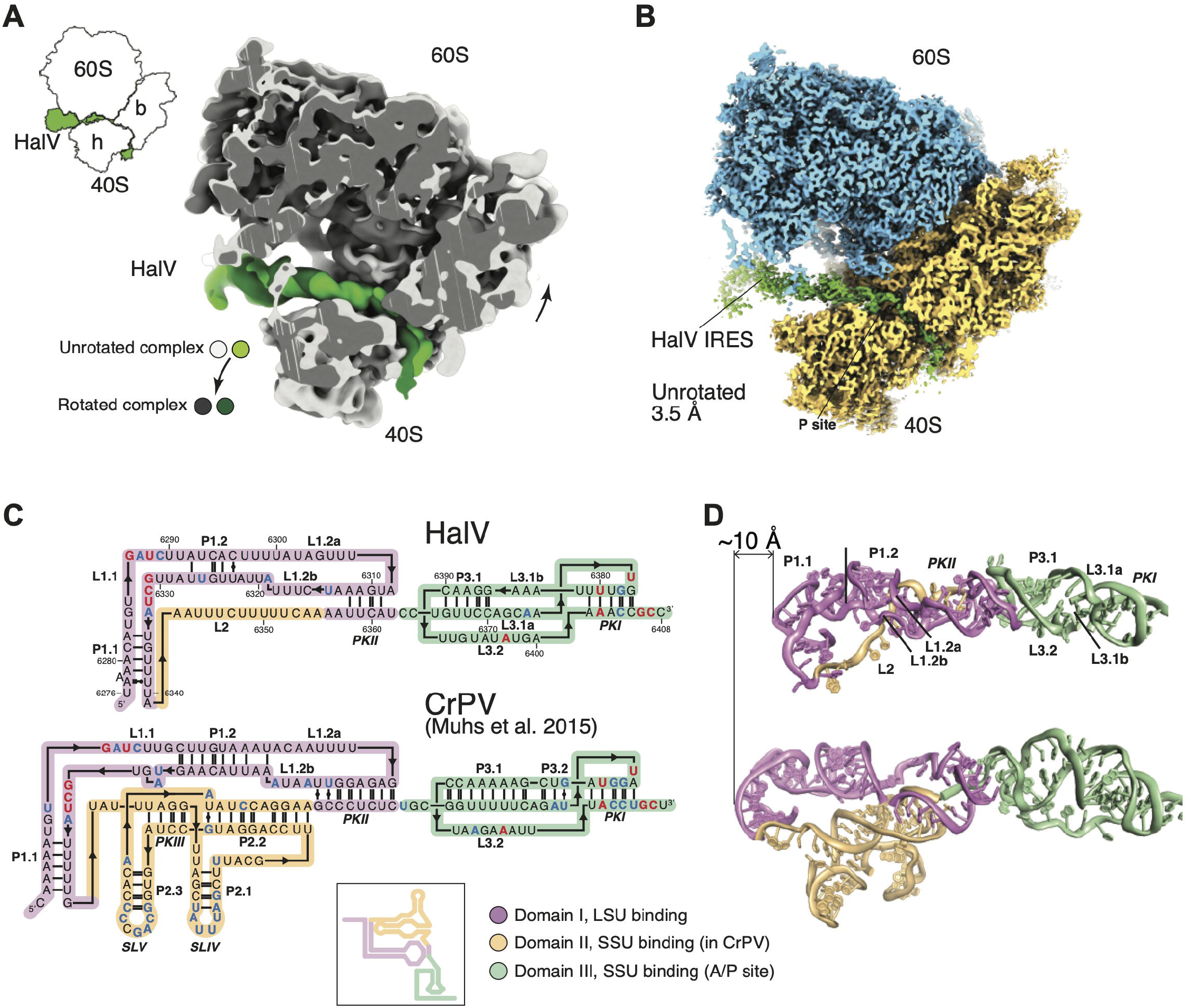
Overview of the HalV IGR IRES bound to the *O. cuniculus* 80S ribosome. (A) Superimposition of the 60S subunit from the unrotated and rotated complexes, to emphasize 40S and IRES movement. (B) Cryo-EM map of the unrotated complex. The 40S is semi-transparent to reveal the path of the IRES. (C) 3D-based secondary structure diagrams of the IGR IRES from HalV (top) and CrPV (bottom). Color coding by domain according to functional role. Residue conservation: >80%, red; 60–80%, blue (within a set of 19 IGR IRES sequences (see Figure S2)). Inset, classical representation of a secondary structure diagram for the CrPV IRES. (D) Comparison of the overall structures of HalV and CrPV. Colorcoding as in panel C.

Modeling of the HalV IGR IRES was based on the predicted secondary structure and tertiary structure elements described above (Figure 3A). The structure of the HalV IGR IRES was mainly built by homology modeling using various previously determined structures of CrPV and PSIV IGR IRESs (see Materials and Methods; Pfingsten et al., 2006; Schüler et al., 2006). The 3D structure of the HalV IGR IRES is similar in the unrotated and rotated states of the IRES/80S complex (Figures S6A-D), and is shorter than the CrPV IGR IRES by ~10 Å (~3 base pairs) (Figure 4D). In sum, the conserved L1.1 loop and PKI (Figures S2 and 4C), which bind to the L1 stalk and to the P site respectively (Figure 4B), are closer to one another in the HalV than in the CrPV IGR IRES (Figure 4D).

The HalV IRES also differs from the CrPV IRES in containing fewer Watson-Crick base-pairs in conserved helical segments, except for the peripheral P1.1 and P3.2 (Figure 4C). Although domain 3 is 60Å long in both IRESs, the longest continuous stack of base-pairs is 11 in CrPV but only 5 in HalV (Figures 4C-D). The atomic model of the unrotated 80S-bound to HalV IRES was fitted (using molecular dynamics flexible fitting, see Methods) into its rotated counterpart. Although the rotated HalV RES/80S map is relatively poorly resolved in the 40S and IRES regions, it was nevertheless possible to analyze the conformational changes of the IRES upon rotation of the 80S ribosome. The implication that the HalV IRES is more flexible was confirmed by the density maps, which revealed a bulge in the central region of the rotated state (Figure 5A). This area of the structure comprises three single stranded regions: L1.2a, L1.2b, and L2. Movement of IRES domain 3 upon subunit rotation (vectors with larger amplitude, Figure 5B) results in a compression of the RNA in that area (vectors with smaller amplitude, Figure 5B). Analysis of the atomic displacement parameters (ADP) indicated that the central region of the IRES, which does not interact with the 80S, is the most dynamic (Figure 5C). Notably, mutations in the HalV-specific L2 loop had relatively low effect on the activity (Figure S3), whereas substitutions in this central region and in the region that binds uL1 were amongst the most deleterious (Figures 3G and 5D).

**Figure 5.**
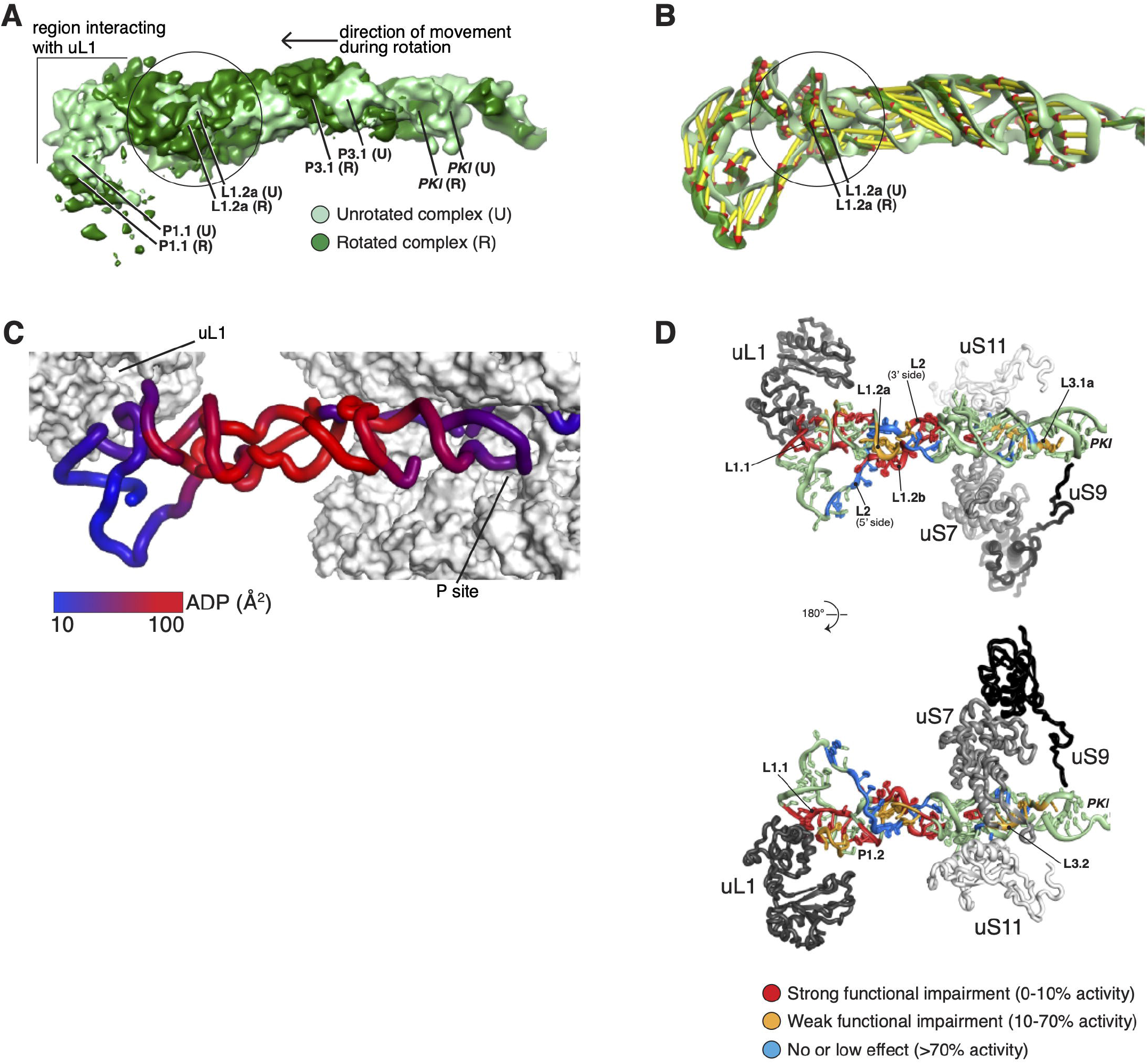
Flexibility of the HalV IRES is key to its function. (A) Compression of the IRES in its central region during rotation. Superimposition of filtered density maps (Gaussian filter 1.5) of the unrotated and rotated states of the 80S-bound IRES. The circle indicates the region where additional bulging density is observed in the rotated complex. (B) Superimposition as in (A) of the IRES 3D models (color-coding as in (A)). Vectors help visualize the direction of the movement, as well as its amplitude. Vectors were calculated by measuring the distance between phosphate atoms. (C) The central region of the IRES is the most dynamic. Color coding of the unrotated IRES by atomic displacement parameter (ADP). (D) Mutations that impair function cluster to regions interacting with ribosomal proteins, and to the central region. Color-coding of the IRES according to the effect of mutations on its function. Percentage of activity in comparison to wild-type RNA.

### The HalV IRES interacts with a subset of the ribosomal proteins bound by canonical IGR IRESs

Interactions with ribosomal proteins occur at the extremities of the HalV IGR IRES, i.e. the conserved L1.1 elbow, and the PKI/loop L3.2 region (Figure 6). The downstream ORF2 also interacts with ribosomal protein uS9 at the mRNA entry site. Protein-RNA interactions mostly involve backbone atoms of the viral RNA and basic amino acids from ribosomal proteins. HalV L1.1 interacts with uL1 similarly to previously characterized IGR IRESs (e.g. Schüler et al., 2006; Fernández et al., 2014; Abeyrathne et al., 2016), i.e. with the a helix of uL1 formed by residues 121-127, as well as loop residues Arg48 (unrotated)/Gln44 (rotated state) and Lys161. The significance of the IRES residues that are predicted to be involved into interaction with uL1 was confirmed by extensive mutagenesis of the L1.1 loop (Figure 3G, left panel). However, as with prior structures of IGR IRESs, precise details concerning the interactions could not be established due to the lower resolution of the map in this area. PKI and L3.2 interact with the ribosomal proteins uS9 (Arg146 interacting with G_6377_), uS7 (Arg135 and Arg136 within interacting distance of nt. 6396-6400), and uS11 (Lys63 and Asp67 near nt. 6395-6397). The latter interaction disappears in the rotated state (compare corresponding panels in Figure 6A and 6C). The predicted interactions are consistent with the inactivating mutations in L3.2 loop and PKI (G_6377_) (Figures 3G, right panel, and 7A). Nt. 6411-6420 in ORF2 bend around uS3, interacting mostly with Arg residues. Many of these interactions are also seen with type VIa and type VIb IGR IRESs (e.g. Abeyrathne et al., 2016; Acosta-Reyes et al., 2019; Muhs et al., 2015), but several of their other characteristic interactions were not observed for the HalV IRES. Thus, although interaction of uL5 with the PKII region is a hallmark of CrPV IRES-80S ribosome complexes (Schüler et al., 2006), the HalV IRES does not come closer than ~18 Å to uL5. Due to the absence of SLIV and SLV-like elements (Figure 4C), the HalV IGR IRES does not interact with eS28 nor with eS25, which remain ~20 Å away in the unrotated state and ~13 Å away in the rotated state.

**Figure 6.**
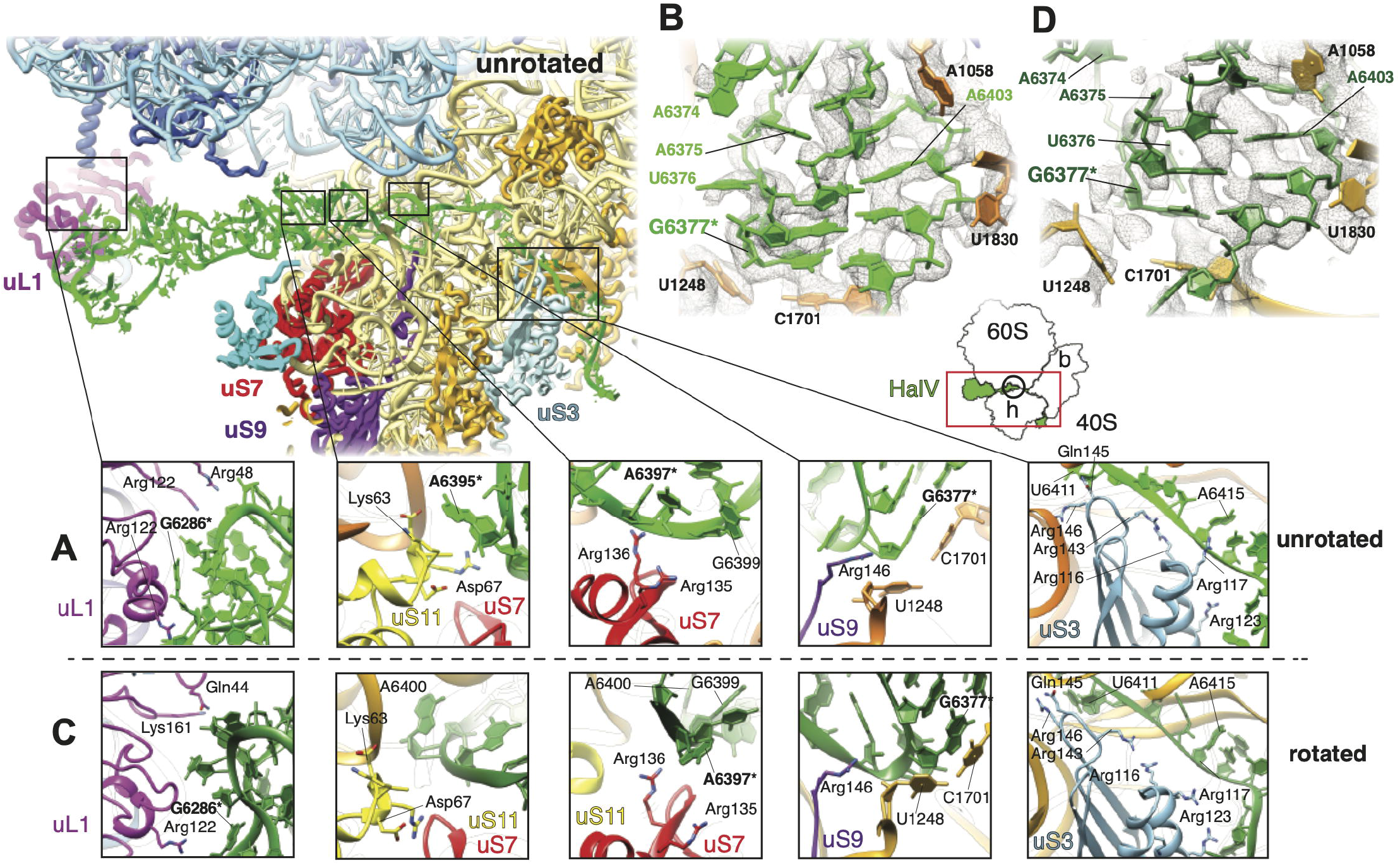
Contacts between the HalV IRES and ribosomal proteins. Overview of a cross section of the ribosome showing the 80S-bound HalV IGR IRES in the unrotated state. The exposed section is indicated on the small 80S/HalV IRES schema on the right by a red rectangle. (A) Blow-up figures on the HalV IRES interaction with the 80S ribosome in the unrotated state. (B) Interactions between PKI and 18S rRNA in the unrotated state. (C) View as in (A) for the rotated state. Note the missing interaction between the HalV IRES and uS11 in this state. (D) View as in (B) for the rotated state. The exposed area is indicated on the small 80S/HalV IRES schema below (B) and (D) by a black circle. IRES nucleotides, which interact with ribosomal proteins and 18S rRNA and which were shown by mutational analysis to be important for IRES function (Figures 3G and 7A), are marked with the asterisks. Note that the PK1 region in the rotated state is more flexible than in the unrotated state, as demonstrated by the scanter local densities and resolution compared to the unrotated state. Color coding for all panels: 40S, dark/light yellow; 60S, dark/light blue; HalV, dark/light green; uL1, magenta; uS11, yellow; uS7, red; uL5, teal; uS3, light blue; uS9, purple.

**Figure 7.**
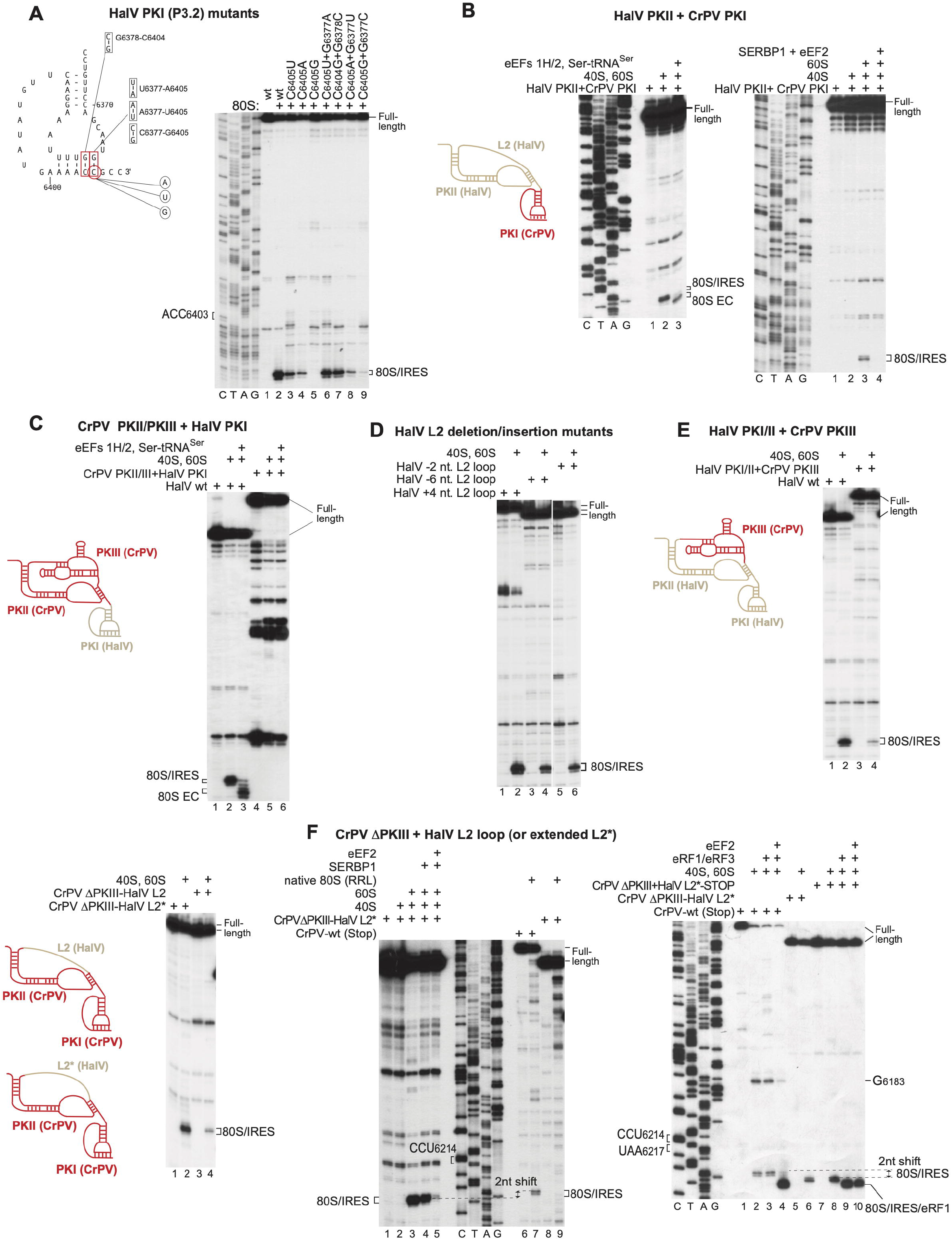
The P site stabilizing interaction of PKI, and activities of hybrid HalV/CrPV IGR IRESs. (A) *Left panel* - Model of the HalV IGR PKI structure, showing nucleotides targeted for substitution to assay a stabilizing interaction of PKI in the P site. *Right panel* – Binding of 80S ribosomes to the indicated HalV IGR IRES mutants, assayed by toe-printing. (B, C, E, F) Ribosomal binding, one-cycle elongation and release factor binding on *wt* and hybrid HalV/CrPV IGR IRESs (shown schematically in each panel) in the presence of the indicated translational components, assayed by toe-printing. The positions of ribosomal complexes are shown. (D) Ribosome-binding activity of the L2 deletion/insertion HalV IGR IRES mutants, assayed by toe-printing. Separation of lanes by white lines indicates that they were juxtaposed from the same gel. (A-F) The positions of ribosomal complexes are indicated. (A, B, F) Lanes *C*, *T, A*, and *G* depict CrPV or HalV sequences.

### PKI of the HalV IGR IRES mimics the tRNA-mRNA interaction in the P site

In the unrotated HalV IGR IRES/80S structure, PKI’s conformation and its interactions with the rRNA and ribosomal proteins in the P site are like those of previously characterized IGR IRESs in posttranslocated complexes (Figures S6E and S7; Costantino et al., 2008; Muhs et al., 2015; Fernández et al., 2014; Zhu et al., 2014; Koh et al., 2014). Thus, G_6377_ in the PKI base-pair adjacent to ORF2 stacks against C_1701_ and interacts with U_1248_ of 18S rRNA (Figure S6E). The importance of these interactions is discussed below. It is interesting to note that in the rotated state, the PKI of the HalV IRES presents scanter densities compared to the more stable unrotated conformation (Figures 6B and 6D).

In the HalV IGR IRES/80S complex in its rotated state, A_1058_ of 18S rRNA interacts in the minor groove of the HalV U_6380_-A_6402_ pair, and U_1830_ contacts C_6404_ (Figure 6D). In this state, the U_6381_-A_6401_ basepairing has been disrupted, so that PKI contains four instead of five base pairs (Figure S7, solid box). The structural changes in PKI are accompanied by conformational changes in L3.2, although the density for the loop residues is too weak to pinpoint actual nucleotide conformations, especially in the rotated state. As suggested for CrPV and TSV IRESs (Ruehle et al., 2015; Abeyrathne et al., 2016), these changes could influence the conformation of domain 3, affecting translocation or stabilizing PKI in the P site. Like other canonical IGR IRESs, the HalV IGR IRES should be able to interact with different ribosomal binding sites during the initiation to elongation transition. Our structures thus illustrate the dynamic nature of PKI in the HalV IGR IRES, which can adjust the number of base pairs it contains that eventually could lead to complete disruption of helix P3.2 on exiting the ribosome as in the CrPV IGR IRES (Pisareva et al., 2018; Figure S7, dashed box).

### Stabilizing interactions of PKI in the P site

The interactions of G_6377_ in PKI with U_1248_ and C_1701_ of 18S rRNA (Figures 6B, 6D and S6E) parallel stabilizing interactions established by CrPV and PSIV PKI (Zhu et al., 2011; Muhs et al., 2015) and stacking of 16S rRNA C_1400_ on the wobble base of the tRNA anticodon and packing of G_966_ against its ribose moiety (Selmer et al., 2006; Korostelev et al., 2006). The observation that compensatory substitutions in P3.2 of PKI failed to fully restore IRES function (Figure 3E) suggests that the interaction of PKI with the P site may impose specific sequence requirements on one or more of the base-pairs in PKI that mimic the initiating codon-anticodon base-pair. This hypothesis was tested by mutational analysis (Figure 7A). Disruption of the G_6377_-C_6405_ base-pair by mutating C_6405_ reduced or even abrogated IRES function (Figure 7A, lanes 3-5). Activity was fully restored by A_6377_-U_6405_ substitutions, but not by U_6377_-A_6405_ or C_6377_-G_6405_ substitutions (Figure 7A, lanes 6, 8 and 9). In contrast, an IGR mutant with a G_6378_-C_6403_→C_6378_-G_6403_ switch in the adjacent base-pair retained nearly full activity (Figure 7A, lane 7). The requirement for a purine at the position of G_6377_ in PKI is consistent with the conservation of base-paired purines at this position in HalV-like and canonical IGR IRESs (Figure S2; Nakashima and Uchiumi, 2008).

### The activities of chimeric HalV/CrPV IGR IRESs

IGR IRESs have a modular organization and it is thus feasible that their evolution involved recombinational exchange of domains. We therefore investigated the activities of various HalV/CrPV IRES hybrids. Replacement of PKI in the GCC_6406_(Ala)→UCU(Ser) HalV IRES by CrPV PKI yielded an IRES that could bind to 80S ribosomes, accept cognate aa-tRNA into the A site and undergo translocation (Figure 7B, left panel). Its binding to 80S ribosomes was abrogated by eEF2/SERBP1 (Figure 7B, right panel), which is characteristic of HalV but not CrPV IRESs (Figures 2C and 2E). Interestingly, the reverse exchange of PKI in the CrPV IRES by HalV PKI did not yield an active hybrid (Figure 7C). Thus, the specific mode of ribosomal binding of CrPV IRES that does not involve initial direct placement of PKI in the P site might not be compatible with the shorter P3.1 stem of HalV PKI.

Next, we investigated the effect of replacement of HalV L2 loop by CrPV PKIII and vice versa. In addition to substitutions (Figure S3), the HalV L2 loop tolerated small insertions and deletions (Figure 7D). However, replacement of the L2 loop by CrPV PKIII yielded a chimeric IRES with a more strongly reduced ribosomal binding activity (Figure 7E). The reverse exchange of CrPV PKIII by nt. 6342-6354 of the HalV L2 loop yielded the CrPVΔPKIII/HalV L2 hybrid that had low ribosome binding activity, but extending the loop to 18 nts (the CrPVΔPKIII/HalV L2* variant) substantially increased the activity of the hybrid IRES (Figure 7F, left panel). This chimeric CrPVΔPKIII/HalV L2* IRES became sensitive to inhibition by SERBP1 and eEF2 like the HalV IRES and also lost the ability to bind to 40S subunits and native 80S ribosomes (Figure 7F, middle panel). Strikingly, it bound directly to the P site leaving the A site accessible without prior eEF2-mediated pseudo-translocation. Thus, 80S ribosomal complexes with the CrPVΔPKIII/HalV L2* IRES yielded toe-prints that were shifted forward by 2nt compared to the wt CrPV IRES (Figure 7F, middle panel, compare lanes 3 and 7, and right panel, compare lanes 2 and 6). Moreover, 80S complexes formed on a variant of this IRES with the A site GCU_6217-6219_→UAA stop codon substitution were able to bind eRF1•eRF3 independently of eEF2, yielding a 2nt toe-print shift like the HalV IRES instead of the *wt* CrPV IRES (Figure 7F, right panel, compare lanes 3-4 and 9-10). A change in the mode of ribosomal binding of the CrPVΔPKIII/HalV L2* IRES was also evidenced by the disappearance of the G_6183_ toe-print (Figure 7, right panel, compare lanes 1-4 with lanes 5-10), which is a hallmark of ribosomal binding of the *wt* CrPV IGR IRES to 80S ribosomes (e.g. Wilson et al., 2000; Pestova and Hellen, 2003; Muhs et al., 2015). Thus, the difference in the ribosomal binding of CrPV and HalV IRESs is determined by the presence of the PKIII domain in the former.

## DISCUSSION

We report that the Halastavi árva virus intergenic region contains an IRES that is related to CrPV-like dicistrovirus IGR IRESs but differs from them structurally and functionally in key respects. Thus, the HalV IGR IRES comprises equivalents of domains 1 and 3 of CrPV-like IRESs, but it lacks the equivalent of domain 2, which enables CrPV-like IRESs to bind stably to the 40S subunit (e.g. Costantino and Kieft, 2005). As a result, the HalV IGR IRES binds directly only to 80S ribosomes. More importantly, PKI of this IRES, which mimics the codon-anticodon interaction, is placed in the ribosomal P site, so that the A site containing the first codon is directly accessible for decoding by eEF1A/aa-tRNA without the prior eEF2-mediated movement of the PKI from the A to P site that is required for initiation on CrPV-like IGR IRESs (Fernández et al., 2014). Thus, the HalV IRES functions by the simplest initiation mechanism known to date: binding of the IRES to 80S ribosomes places the [codon-anticodon]-mimicking PKI directly into the ribosomal P site, which is followed by eEF1A-mediated decoding of the A site codon.

The elements of the HalV IGR IRES that interact with the 80S ribosome are restricted to the IRES’s extremities. Thus, the L1.1 loop contains conserved motifs like those in CrPV-like IGR IRESs and, analogously, interacts with ribosomal protein uL1 in the L1 stalk of the 60S subunit, whereas at the opposite end, elements in PKI interact with protein and rRNA constituents of the ribosomal P site. Binding of PKI in the P site is stabilized by G_6377_, which interacts directly with universally conserved elements of the P site, and by the L3.2 loop, as has been suggested for the post-translocation canonical IGR IRES (Abeyrathne et al., 2016). Consistent with direct binding to the P site, the HalV IGR IRES is ~10Å shorter than CrPV-like IGR IRESs, reflecting that the P site is closer than the A site to the L1 stalk. The HalV IGR IRES is also intrinsically less ordered, with fewer base-pairs in most conserved helical segments, and more flexible than canonical IGR IRESs. The higher flexibility was already apparent from the density maps, which revealed a bulge in the central region of the rotated state of the IRES, an area that comprises three single-stranded regions: L1.2a, L1.2b, and L2. The flexibility appears to be important for IRES function, because substitutions in L1.2a and L1.2b strongly reduced IRES function, potentially by introducing stabilizing base-pairs that could increase rigidity. However, the biggest difference between the HalV IRES and previously characterized IGR IRESs is the lack in the former of the equivalent of domain 2, which contains the conserved stem-loops IV and V that interact with uS7 and eS25 in the 40S subunit (e.g. Schüler et al., 2006; Abeyrathne et al., 2016). Although it is well established that domain 2 enables stable binding of type VIa and type VIb IGR IRESs to 40S subunits (Jan and Sarnow, 2002; Nishiyama et al., 2003; Costantino and Kieft, 2005), we found that it is also responsible for positioning the PKI outside the P site. Thus, deletion of domain 2 from the CrPV IGR IRES resulted in direct binding of PKI to the P site, which eliminated the necessity for the initial step of eEF2-dependent translocation of PKI of the IRES. In the absence of domain 2, even the greater length of the CrPV IGR IRES did not induce slippage of PKI into the A site. Notably, lengthening the linker that replaced domain 2 in the CrPV IGR IRESΔPKIII mutant increased its efficiency of action, likely reflecting the importance of the IRES flexibility for the new mode of ribosomal binding.

Importantly, we observed that ribosomal binding of the HalV IGR IRES is inhibited by SERBP1/eEF2, which associate with non-programmed eukaryotic 80S ribosomes (e.g. Anger et al., 2013; Zinoviev et al. 2015; Brown et al., 2018). As a result, the translational efficiency of the HalV IRES in RRL was very low. In contrast, the *wt* CrPV IGR IRES was resistant to SERBP1/eEF2, but deletion of domain 2 rendered the CrPV IGR IRES susceptible to such inhibition. Thus, the presence of domain 2 on an IGR IRES not only determines the ribosomal position of PKI, but also confers resistance to SERBP1/eEF2 inhibition. This raises the question of how the HalV IGR IRES overcomes such inhibition *in vivo*. HalV was derived from the intestinal content of carp, and one possibility is that its natural host, which has not yet been established, may lack SERBP1 or encodes a homolog that does not impair translation in this way. Another possibility is that the viral infection could influence the cytoplasmic conditions, leading to relief of this inhibition. In this respect it is worth mentioning that although initial studies had suggested that domain 2 contains elements that are critical for the function of canonical IGR IRESs (e.g. Jan and Sarnow, 2002), dicistrovirus infection is now known to alter the intracellular environment in a manner that suppresses the defect in IRES function caused by mutations in this domain (Kerr et al., 2016). The nature of this change has not been established, but it is possible that stress-induced changes in the localization of SERBP1 (e.g. Lee et al., 2014), its dissociation from ribosomes or even its degradation during infection could lead to activation of IRES function. In this context it would be of interest to determine the potential influence of other 80S-binding proteins such as IFRD2 and CCDC124 (Brown et al., 2018; Wells et al., 2020) on the activity of HalV and canonical IGR IRESs.

Our metagenomic analyses determined that HalV is only one of a family of related IGR elements that lack an equivalent of domain 2. Although we cannot exclude the possibility that they derive from canonical IGR IRES progenitors as a result of loss of a functional element (i.e. domain 3), a more likely scenario is that HalV-like IRESs form an ancestral class (VIc). According to that model, the presence of domain 3 in type VIa and type VIb IRESs would represent an evolutionary acquisition, favored by a resulting gain of function. Perhaps this gain was the switch to initiation from P to A sites, which has been suggested to facilitate post-binding events by partitioning the energetic penalty for the required dissociation of the IRES after initial recruitment to the ribosome (Muhs et al., 2015). It could also have been resistance to inhibition by SERBP1/eEF2 or other ribosome-binding proteins. In this model, IRES evolution involves accretion of domains, as suggested for the evolution of ribosomal RNA (Petrov et al., 2014) and hepatitis C virus-like IRESs (Asnani et al., 2014) and their subsequent dissemination to other viral genomes by recombination. The number of novel, often divergent dicistroviruses is increasing rapidly (e.g. Shi et al., 2016) and it will be interesting to determine whether there are IGR IRESs that lack domain 2 but nevertheless still bind stably to 40S subunits or that contain some equivalent of domain 2 but nevertheless bind in the P site. An exciting possibility is that IGR-like elements with divergent domain 1 or domain 2 elements may have evolved the ability to exploit other potential sites of interaction, leading to utilization of divergent mechanisms of initiation.

## ACKNOWLEDGEMENTS

We thank Cajetan Neubauer and Venki Ramakrishnan for their gift of RelE, and Alexander Myasnikov for assistance in data acquisition. This work was supported by NIH grant GM122602 to T.V.P., NIH grant AI123406 to C.U.T.H., and by the ANR grants ANR-14-ACHN-0024 @RAction program ‘‘ANR CryoEM80S’’, ANR-11-LABX-0057_NETARN and the ERC-2017-STG #759120 “TransTryp” to Y.H.. This project was supported by “Institut National de la Santé et de la Recherche Médicale” (INSERM), “Centre National de la Recherche Scientifique” (CNRS) and Université de Bordeaux.

## AUTHOR CONTRIBUTIONS

ISA conducted all biochemical experiments. ISA, TVP and CUTH interpreted biochemical data. CUTH conducted bioinformatic and phylogenetic analysis. AS made the cryo-EM grids and supervised data-acquisition. HS and YH processed the cryo-EM data. QV, AB and YH built the atomic models and interpreted the structures. ISA, QV, TVP, YH and CUTH wrote the manuscript. TVP, YH and CUTH directed the research.

## KEY RESOURCES TABLE

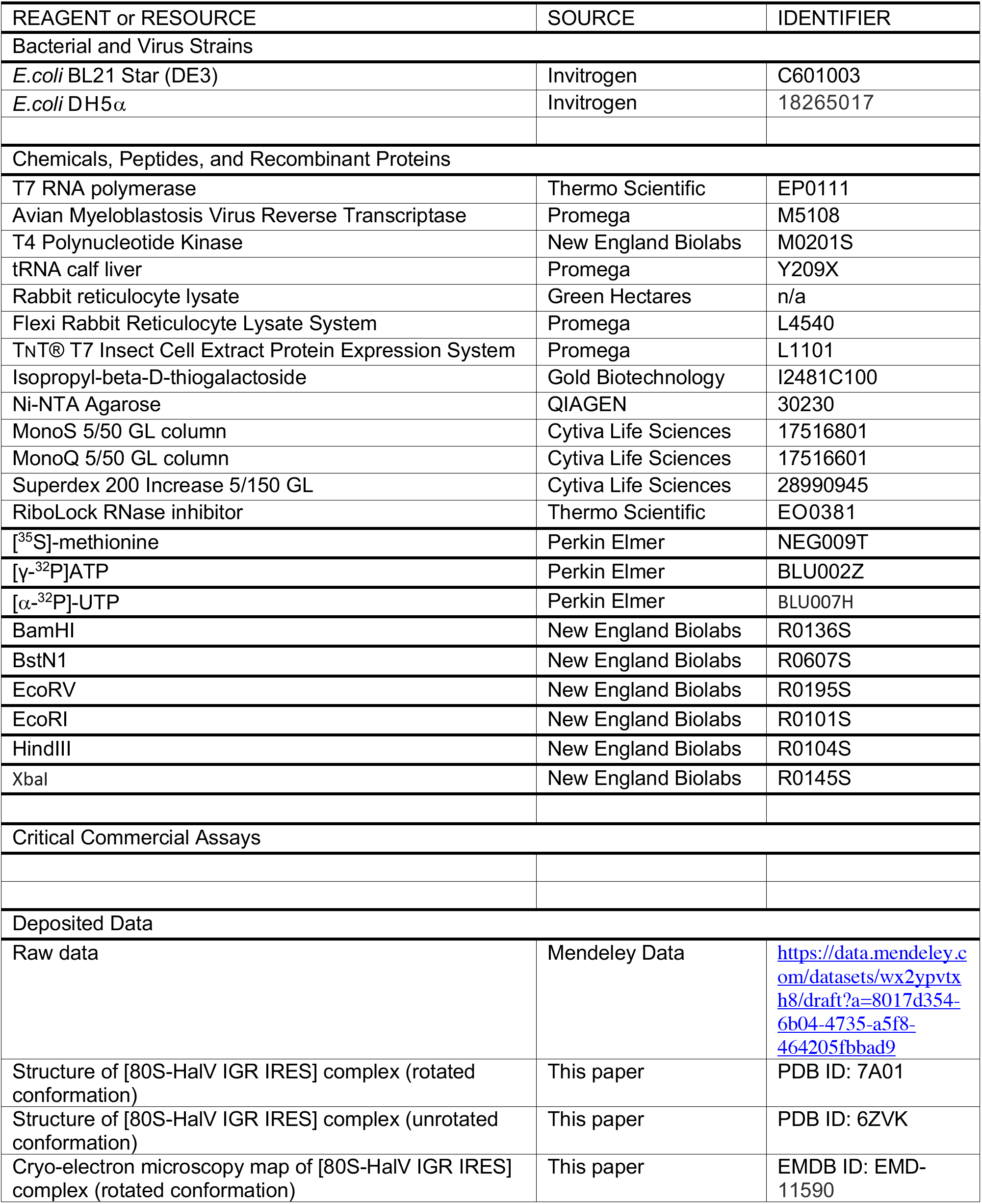
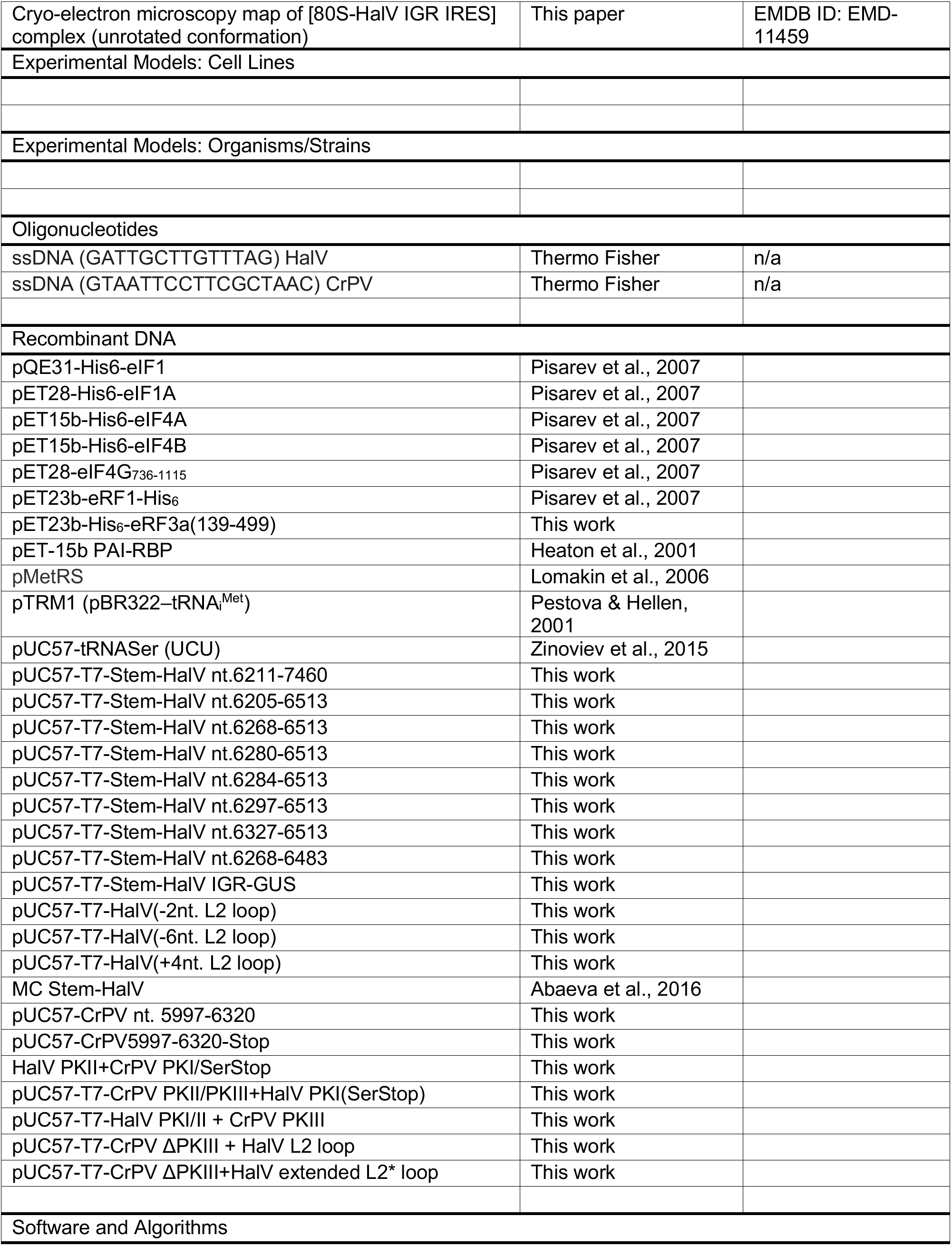
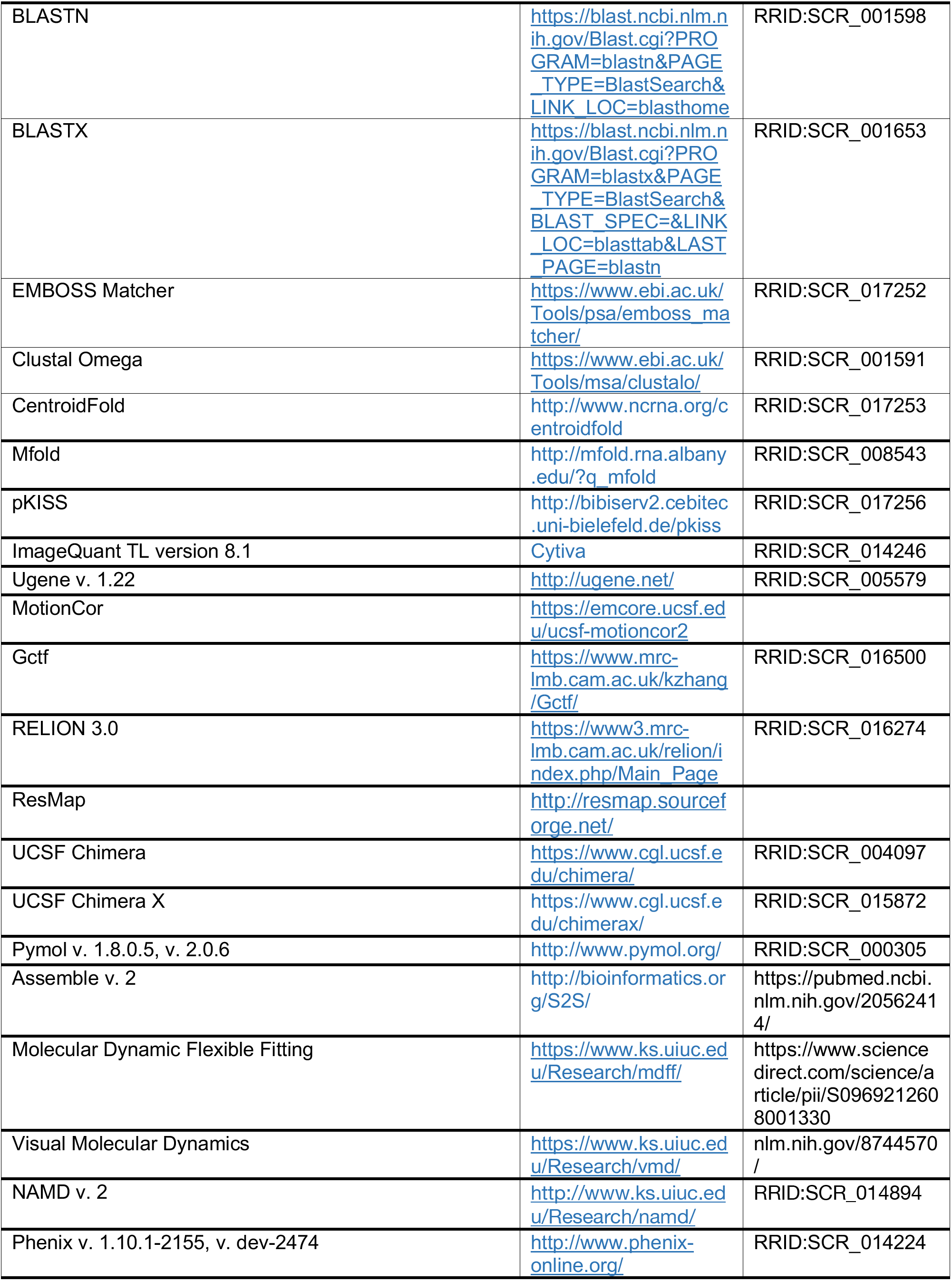
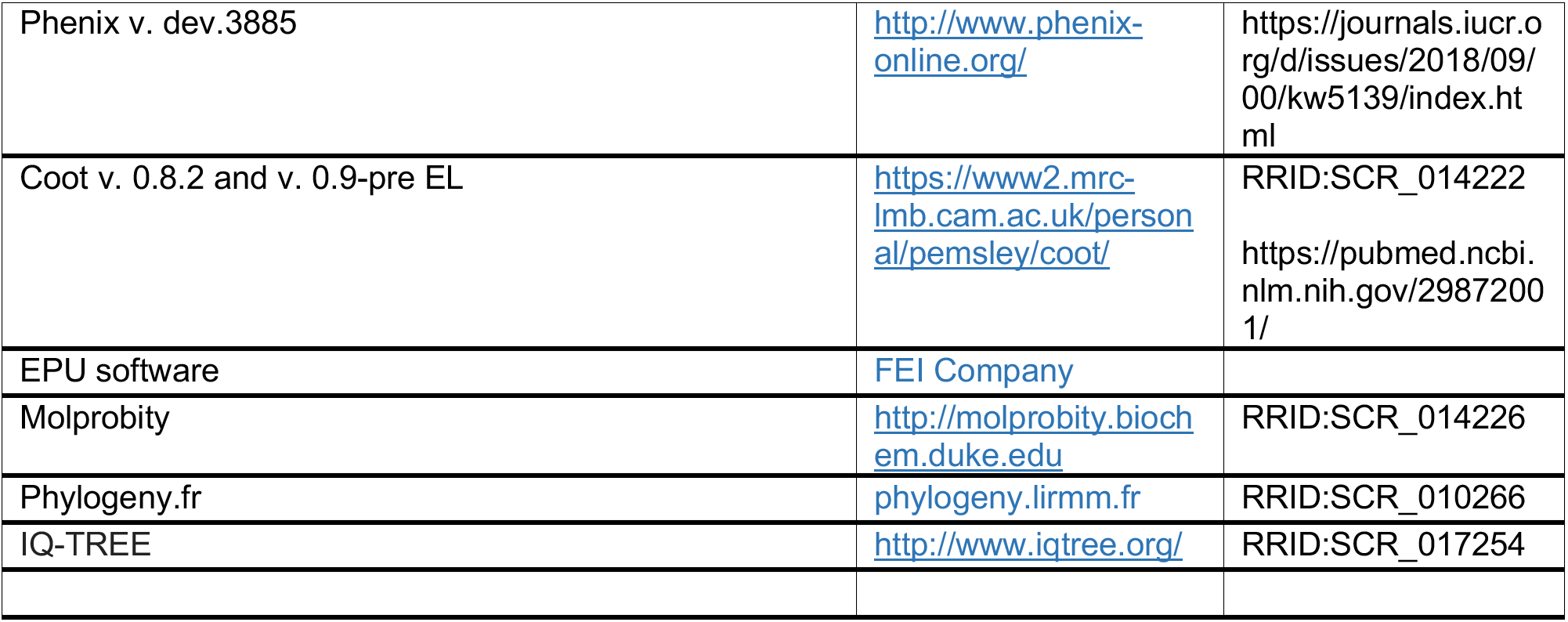

## LEAD CONTACT AND MATERIALS AVAILABILITY

Further information and requests for resources and reagents should be directed to and will be fulfilled by the Lead Contact, Tatyana Pestova. All unique/stable reagents generated in this study are available from the Lead Contact without restriction.

## EXPERIMENTAL MODEL AND SUBJECT DETAILS

Plasmids were propagated in *Escherichia coli* DH5a cells and recombinant proteins were expressed in *E. coli* BL21(DE3) cells, respectively, both grown in LB medium.

## METHOD DETAILS

### Construction of Plasmids

Vectors for expression of His_6_-tagged eIF1 and eIF1A (Pestova et al., 1998a), eIF4A and eIF4B (Pestova et al., 1996), eIF4G_736-1115_ (Lomakin et al., 2000), *Escherichia coli* methionyl tRNA synthetase (Lomakin et al., 2006), Serbp1 (Heaton et al., 2001) and eRF1 (Seit-Nebi et al., 2001) have been described. A vector for expression of His_6_-tagged eRF3a (a.a. 139-499) lacking the N-terminal 138 a.a. (referred to as eRF3 in the text) was made by inserting DNA between BglII and Nde1 sites of pET23(b) (GenScript, Piscataway, NJ).

The transcription vectors for tRNA^Ser^ (UCU codon) (Zinoviev et al., 2015) has been described. New transcription vectors were made by inserting DNA into pUC57 (GeneWiz, South Plainfield, NJ) or by subsequent site-directed mutagenesis (NorClone Biotech, London, ON, Canada; 96 Proteins, San Francisco, CA). The vector monocistronic Stem-HalV nt.6211-7460 contained DNA corresponding to a T7 promoter, a stable hairpin (GGCCGACCCGGTGACGGGTCGGCC) (ΔG=-25.80 kcal/mol), HalV nt. 6211-7460 and a linker comprising AvrII, SmaI, ClaI and XbaI sites. The vectors for truncated HalV IGR IRES mutants contained a T7 promoter followed by the corresponding HalV sequences (nt.6205-6513, 6268-6513, 6280-6513, 6284-6513, 6297-6513 and 6327-6513) and restriction sites for EcoRV and EcoRI. The vectors for HalV IGR IRES substitution/deletion/insertion mutants were made using a base construct containing nt. 6268-6483.

The transcription vector Stem-HalV IGR-GUS (employed for synthesis of mRNA used for translation in rabbit reticulocyte lysate (RRL)) included a T7 promoter, the hairpin described above, HalV nt.6268–6513 (with a GCC_6406-8_UCU substitution, as well as a AUG_6397-9_UAG substitution that eliminates a possibility of detecting a product originated from non-specific AUG-dependent initiation within the IRES), a modified fragment of the *E. coli* beta-glucuronidase (GUS) ORF (GenBank: AAA68923.1), HalV nt. 7404-7460 and an [AvrII-Sma1-Cla1-Xba1] linker. The transcribed mRNA encodes a 286 a.a.-long (31.7 kDa) fusion protein comprising a Serine residue and a.a. 2-36 of the HalV ORF2 linked to a modified form of a.a. 351-602 of the GUS ORF in which AUG codons 407 and 446 had been substituted and codons 528, 548, 560, 569, 583 and 596 had been replaced by AUG triplets. The MC Stem-HalV transcription vector (employed for synthesis of mRNA used for translation in RRL) containing the HalV 5’UTR and an adjacent 256 a.a.-long ΔORF1 (nt. 1-1682) has been described (Abaeva et al., 2016).

Vectors for transcription of CrPV IGR IRES mRNAs contained CrPV nt. 5997-6320 or a variant (CrPV_5997-6320_-Stop) with GCT_6217-9_TAA substitutions inserted between BamH1 and EcoR1 sites of pUC57 (GeneWiz). The vectors HalV PKII+CrPV PKI/SerStop and CrPV PKII/PKIII+HalV PKI(SerStop) for transcription of hybrid HalV-CrPV IGR IRESs contained HalV nt.6268-6365 + CrPV nt.6174-6320 (GCTACA_6217-6222_→TCTTAA) and CrPV nt. 5997-6173 + HalV nt. 6366-6483(GCCACT_6406-6411_→TCTTAA) downstream of T7 promoters, respectively. The vectors for transcription of HalV L2 −2 nt. L2 loop, HalV −6 nt L2 loop and HalV +4 nt. L2 loop mutant mRNAs were made by replacing the L2 loop sequence AAUUUCUUUUUCAA by the sequences AAUUCUUUUCAA, AAUCUCAA and AAUUUUUCUUUUUUUCAA, respectively. The vector for transcription of HalV PKI/II + CrPV PKIII mRNA was made by replacing HalV nt. 6342-6353 inclusive by CrPV nt. 6110-6162. The transcription vectors CrPV ΔPKIII + HalV L2 loop and CrPV ΔPKIII + HalV extended L2* loop were made by replacing CrPV nt. 6110-6161 by HalV nt. 6343-6353 and by the HalV-related sequence AAUUUUUCUUUUUUUUC, respectively.

All RNAs were transcribed using T7 RNA polymerase (Thermo Scientific).

### Purification of factors, ribosomal subunits and aminoacylation of tRNA

Native mammalian 40S and 60S subunits, eIF2, eIF3, eEF1H, eEF2 and total aminoacyl-tRNA synthetases were purified from rabbit reticulocyte lysate (RRL), and insect 40S and 60S subunits were purified from *S. frugiperda* cell-free extract (Promega) as described (Pisarev et al., 2007). Native 80S ribosomes were purified from 200 μl RRL (Promega), which was layered onto 10-30% sucrose density gradients in buffer A (20 mM Tris [pH 7.5], 100 mM KAc, 2.5 mM MgCl_2_, 2 mM DTT and 0.25 mM spermidine) and centrifuged in a Beckman SW55 rotor at 53,000 rpm for 90 min at 4°C. Fractions were collected across the gradient, and the position of 80S ribosomes was determined by monitoring the absorbance at 260 nm. Recombinant eIF1, eIF1A, eIF4A, eIF4B, eIF4G_736-1115_, eRF1, eRF3, *Escherichia coli* methionyl tRNA synthetase and SERBP1 were expressed and purified from *E. coli* as described (Pisarev et al., 2007; Alkalaeva et al., 2006; Zinoviev et al., 2015). RelE was a gift from Venki Ramakrishnan. Native total calf liver tRNA (Promega) and *in vitro* transcribed tRNA^Ser^ were aminoacylated using *Escherichia coli* methionyl tRNA synthetase (for obtaining Met-tRNA_i_^Met^) or total aminoacyl-tRNA synthetases (for aminoacylation of elongator tRNAs) as described (Pisarev et al., 2007; Zinoviev et al., 2015).

### Assembly and analysis of ribosomal complexes by toe-printing

For assembly of 48S initiation complexes, 2 pmol Stem-(*wt* HalV IGR) mRNA, 3 pmol 40S subunits, 6 pmol eIF2, 4.5 pmol eIF3, 8 pmol eIF1, 8 pmol eIF1A, 10 pmol eIF4A, 5 pmol eIF4B, 5 pmol eIF4G_736-1115_ and total tRNA containing 7 pmol Met-tRNA_i_^Met^ were incubated in 40 μl buffer A supplemented with 1 mM ATP and 0.1 mM GTP for 10 min at 37°C. For studying ribosomal asociation of HalV, CrPV and hybrid HalV/CrPV IGR IRESs, 2 pmol mRNA was incubated with 5 pmol 40S subunits, 8 pmol 60S subunits or both in the presence or absence of indicated combinations of 10 pmol eRF1, 10 pmol eRF3, 10 pmol eEF2 and 10 pmol SERBP1 in 40 μl buffer A supplemented with 1 mM ATP and 0.1 mM GTP for 10 min at 37°C. To examine the elongation competence of assembled 80S/IRES complexes, reaction mixtures were supplemented with combinations of 10 pmol eEF2, 10 pmol eEF1H, 500 μg/mL of cycloheximide, 15 μg appropriately aminoacylated native total tRNA or 6 pmol Ser-tRNA^Ser^, and incubation was continued for 10 min at 37°C. Resulting ribosomal complexes were analyzed by toe-printing as described (Pisarev et al., 2007) using avian myeloblastosis virus reverse transcriptase (AMV RT) (Promega) and [^32^P]-labeled oligonucleotide primers complementary to HalV nt. 6458-72 or CrPV nt. 6304-19, as appropriate. cDNA products were resolved in 6% polyacrylamide sequencing gels followed by autoradiography or phosphoimager analysis.

### Assembly and analysis of ribosomal complexes by sucrose density gradient centrifugation

Ribosomal complexes were formed by incubating 10 pmol [^32^P]-labeled HalV IGR IRES mRNA with 30 pmol 40S subunits, 40 pmol 60S subunits or both, as indicated, in buffer A supplemented wth 1 mM ATP and 0.1 mM GTP for 10 min at 37°C, and then resolved by centrifugation through 10-30% sucrose density gradients in buffer A in a Beckman SW55 rotor at 53,000 rpm for 90 min at 4°C. The optical density of fractionated gradients was measured at 260 nm, and the presence of [^32^P]-labeled mRNA was monitored by Cherenkov counting.

### Assembly and analysis of ribosomal complexes by RelE cleavage

Analysis of cleavage of ribosome-bound mRNA by RelE was done as described (Skabkin et al., 2013). 80S ribosomal complexes were assembled in the presence or absence of eEF2 as described above for preparation of complexes for toe-printing analysis, and then incubated with 20 pmol RelE for 10 min at 37°C. After that, mRNA was phenol-extracted and analyzed by primer extension using AMV RT and the same [^32^P]-labeled primers, as described above for toe-printing analysis. cDNA products were resolved in 6% polyacrylamide sequencing gels followed by autoradiography or phosphoimager analysis.

### Viral nucleotide sequences

Sequences were retrieved from the NCBI database (http://www.ncbi.nlm.nih.gov/nuccore) using the following accession numbers: Acute bee paralysis virus (ABPV), NC_002548.1; Aphid lethal paralysis virus 1 (ALPV), NC_004365.1; Beihai mantis shrimp virus 5 (NC_032434.1); Black queen-cell virus (BQCV) (NC_003784.1); Changjiang picorna-like virus 14 (NC_032773.1); Cricket paralysis virus (CrPV), NC_003924.1; Drellivirus strain 93C3 (KX924637); *Drosophila* C virus (DCV) strain EB (NC_001834.1); Empeyrat virus (KU754505.1); Formica exsecta virus 1 (NC_023021.1); Goose dicistrovirus (NC_029052.1); *Halastavi árva* RNA virus (NC_016418.1); Himetobi P virus (HiPV) (NC_003782.1); Homalodisca coagulata virus 1 (NC_008029.1); Israel acute paralysis virus (IAPV), NC_009025.1; Kashmir bee virus (KBV) (NC_004807.1); Kuiper virus (KX657785.1); *Macrobrachium rosenbergii* Taihu virus strain cn-taihu100401(NC_018570.2; Mud crab virus (NC_014793.1); *Nilaparvata lugens* C virus (NlCV), KM270560.1; *Plautia stali* intestine virus (PSIV) (NC_003779.1); *Rhopalosiphum padi* virus (RhPV) (NC_001874.1); Shahe arthropod virus 1 strains SHWC01c3692 (KX883988), SHWCII5326 (KX883671), SHWC0209c12762 (NC_032422) and SHWC0209c12762 (KX883641); *Solenopsis invicta* virus 1 (SinV), NC_006559.1; Taura syndrome virus (TSV), NC_003005.1; *Triatoma* virus (TriV) (NC_003783.1); Wenzhou shrimp virus 7 (NC_032420.1) and the TSA Sequence *Proasellus solanasi* HAFJ01006557.1.

### Phylogenetic analysis (Figure S1D)

Dicistrovirus ORF2 capsid protein precursor amino acid sequences were obtained by conceptual translation of viral sequences downstream of the IGR IRES, submitted to the web server phylogeny.lirmm.fr (Dereeper et al., 2008) for alignment using CLUSTAL-W (default parameters) and elimination of positions containing gaps, and then used for inference of the maximum likelihood (ML) phylogenetic tree using IQ-TREE (Nguyen et al., 2015), applying the WAG+FO model for amino acid substitution and using 10,000 ultrafast bootstraps (Hoang et al., 2018). Statistical support for individual nodes was estimated using the bootstrap value.

### Identification of candidate IGR IRESs

Candidate IGR sequences were identified using BLASTN (http://www.ncbi.nlm.nih.gov/BLAST/) searches of nucleotide collection (nr/nt) and TSA sequences, and BLASTX searches of non-redundant protein (nr) and Transcriptome Shotgun Assembly (TSA) protein (tsa_nr) sequences in the NCBI database. Nucleotide searches used the parameters: E, 1000; word size, 11; match/mismatch scores, 1/1; gap costs, 2/1, and polypeptide searches used the parameters: E, 1000; word size, 6; Matrix: BLOSUM62; gap costs, 9/1. TSA searches were limited to *Arthropoda* and *Mollusca*, and ‘hits’ were characterized by 6-frame translation. Polypeptide sequences were used in BLAST searches to verify that the C-terminal region of ORF1 encoded the 3D polymerase and that ORF2 encoded capsid proteins. IGR sequences were aligned using EMBOSS Matcher (http://www.ebi.ac.uk/Tools/psa/emboss_matcher/nucleotide.html) and Clustal Omega. Viral 3CD and P1 capsid protein sequences were aligned using Clustal Omega with default parameters to determine pairwise sequence identities.

### In vitro translation (Figure 2A)

MC Stem-HalV and GUS-(HalV IGR) mRNAs (3 pmol) were translated in 20μl reaction volume of Flexi rabbit reticulocyte lysate (Promega) in the presence of [^35^S]methionine (>37.0 TBq/mmol; Perkin Elmer) for 30 minutes at 30°C. When indicated, reaction mixtures were supplemented with 6 pmol 40S and 8 pmol 60S subunits with or without prior preincubation with mRNAs for 5 minutes at 37°C in buffer A supplemented with 1 mM ATP and 0.1 mM GTP. Translation products were resolved by electrophoresis using NuPAGE 4–12% Bis-Tris precast gels (Invitrogen), followed by autoradiography.

### Chemical and enzymatic probing (Figure S1A - S1C)

HalV IGR-containing mRNA was enzymatically digested with RNase V1 or chemically modified with 1-cyclohexyl-(2-morpholinoethyl) carbodiimide metho-p-toluene sulphonate (CMCT), dimethylsulfate (DMS) or *N*-methylisatoic anhydride (NMIA) exactly as described (Abaeva et al., 2016). Cleaved or modified sites were identified by primer extension, using AMV RT and primers complementary to HalV nt. 6353-70 or nt. 6458-72. cDNA products were resolved in 6% polyacrylamide sequencing gels followed by autoradiography.

### Nucleotide sequence alignment (Figure S2)

IGR sequences were aligned automatically using Clustal Omega (http://www.ebi.ac.uk/Tools/msa/clustalo/) and then manually using Ugene v. 1.22 (Okonechnikov et al., 2012), relying on published sequence alignments as a guide (Jan, 2006; Kanamori and Nakashima, 2001; Pfingsten et al., 2006).

### Initial HalV IGR structure modeling

Structural elements were initially modeled using CentroidFold (http://www.ncrna.org/centroidfold) (Sato et al., 2009) and Mfold (http://mfold.rna.albany.edu/?q_mfold) (Zuker, 2003). Tertiary structures were modeled using pKiss (http://bibiserv2.cebitec.uni-bielefeld.de/pkiss) (Janssen and Giegerich, 2015), in all instances using default parameters.

### Grid preparation

15 pmol HalV mRNA (nt. 6268-6483) was incubated with 7.5 pmol 40S subunits and 7.5 pmol 60S subunits in 40 μl buffer B (20 mM Tris [pH 7.5], 100 mM KAc, 5 mM MgCl_2_, 1 mM DTT, 0.25 mM spermidine, 0.5 mM ATP and 0.5 mM GTP) for 5 minutes at 37°C to allow 80S/IRES complex formation. After incubation, the reaction mixture was diluted using buffer B to achieve the concentration of 80S ribosomes of 70 nM, and 4 *μ*L of the sample was applied onto the Quantifoil R2/2 400-mesh holey carbon grid, which had been coated with thin carbon film and glow-discharged. The sample was incubated on the grid for 30 sec and then blotted with filter paper for 1.5 sec in a temperature and humidity controlled Vitrobot Mark IV (T = 4°C, humidity 100%, blot force 5) followed by vitrification in liquid ethane.

### Image acquisition

Data collection was performed on a spherical aberration corrected Titan Krios S-FEG instrument (FEI Company) at 300 kV using the EPU software (FEI Company) for automated data acquisition. Data were collected at a nominal underfocus of −0.6 to −4.5 *μ*m at a magnification of 127,272 X yielding a pixel size of 1.1 Å. Micrographs were recorded as a movie stack on a Falcon II direct electron detector (FEI Company), each movie stack were fractionated into 17 frames for a total exposure of 1 sec corresponding to an electron dose of 60 ē/Å^2^.

### Image processing

Drift and gain correction and dose weighting were performed using MotionCor2 (Zheng et al., 2017). A dose weighted average image of the whole stack was used to determine the contrast transfer function with the software Gctf (Zhang, 2016).The following process was done using RELION 3.0 (Zivanov et al., 2018). Particles were picked using a Laplacian of gaussian function (min diameter 300 Å, max diameter 320 Å). 337,268 particles were extracted with a box size of 360 pixels and binned three-fold for 3D classification into 5 classes. Two sets of subclasses depicting high-resolution features were selected for refinement, one “unrotated” (55589 particles) and one “rotated” (42135 particles). Refinement of unrotated and rotated classes yielded an average resolution of 3.6 Å and 3.5 Å, respectively. The unrotated class has been focused refined with a mask on the 60S, the body and the head of the 40S, yielding respectively 3.49, 3.52 and 4.13 Å resolution. Determination of the local resolution of the final density map was performed using ResMap (Kucukelbir et al., 2014).

### Map fitting and model refinement

A cryo-EM structure of the 80S ribosome from *Oryctolagus cuniculus* (PDB ID 4UJE; (Budkevich et al., 2014)) was fitted in the density map using Chimera v. 1.10.2 (Pettersen et al., 2004). RNA regions that were not seen in this reference structure were built within Assemble v. 2 (Jossinet et al., 2010). Molecular dynamics flexible fitting (MDFF; (Trabuco et al., 2008)) with explicit solvent was performed for the complete 80S ribosome in VMD v. 1.9.2 (Humphrey et al., 1996), using NAMD v. 2 (gscale 0.3, numsteps 500,000, minsteps 2,000) (Phillips et al., 2005). RNA geometry and fit in density were improved by running Erraser within Phenix v. 1.10.1-2155 (Adams et al., 2010; Afonine et al., 2013; Chou et al., 2013; Jain et al., 2015), for rRNA fragments of ~ 990 nt. This model was further minimized using NAMD.

The IGR IRES from HalV within the unrotated complex was principally built by homology with various CrPV and PSIV IRES structures (earlier 3D models based on only partially correct 2D models had been built *ab initio*). More specifically, the IGR IRES was assembled from the following modules: *ab initio* modeled PKI and L3.2; P3.1 and L3.1 from residues 113-120 and 201-207 in PDB ID 1HR2 (double helical region of the P4-P6 domain from the *T. thermophila* group I intron that contains a tandem of purine-purine base pairs (Juneau et al., 2001)); PKII and P1.2 from PDB ID 2IL9 (X-ray crystal structure of the PSIV IRES lacking PKI at 3.1 Å resolution (Pfingsten et al., 2006)); P1.1 and L1.1 from PDB ID 2NOQ (first cryo-EM reconstruction of a complete 80S-bound CrPV IRES (Schüler et al., 2006)). P1.2 required the most attention during manual rebuilding as it is shorter and less rich in Watson-Crick pairs than its counterpart in PSIV.

These modules were connected to one another and placed in filtered density maps (using Gaussian filters with 1.5 and 2.0 widths), using the ‘fit in map’ option in Chimera v. 1.10.2 (Pettersen et al., 2004), and the ‘real space refine’ and ‘regularize zone’ options in Coot v. 0.8.2 (Emsley and Cowtan, 2004). For real-space refinement and subsequent manual rebuilding, the IRES was stripped off of the 80S ribosome, except for RNA and protein residues within ~15 Å of any IRES residue. Upon assembling this partial model, the path for the L2 single strand that replaces the larger SSU-binding domain in CrPV and PSIV became straightforward. This 13-nucleotide long U and C-rich RNA segment wraps around L1.2a and L1.2b and could be modeled using the first residues of the corresponding domain in the CrPV IRES, that were extended using the ‘add terminal residue’ functionality in Coot with HalV-specific residues, in order to connect to PKII. The sequence of the IRES was manually edited using the ‘swapna’ command in Chimera and the ‘simple mutate’ functionality in Coot.

Manual fitting with geometry correction was carried out throughout the entire IRES, with a particular attention to the L1.1/L1 stalk interface, the central region of the IRES that comprises L1.2a, L1.2b, and L2, the L3.2 joining region that interacts with protein eS7, as well as the two pseudoknots. Assigning the correct sequence register for the IRES was made possible by first building the PKI region, which is the most conserved across IGR IRESs (Figures S2 and S6F), and where the density allows purines to be distinguished from pyrimidines (Figure S6). The resulting model was used as input for real-space refinement in Phenix v. dev-2474 (Adams et al., 2010; Afonine et al., 2013), using the unfiltered density map. The refinement procedure included simulated annealing (starting temperature = 600 K) and global minimization for 5-10 macro-cycles and took into account RNA and protein secondary structure restraints (search_method = from_ca).

A close-to-final model of the HalV IRES in the unrotated state was used as the starting model for the IRES in the rotated state. The rotated IRES structure was refined using the same procedure as the unrotated state. Both structures were placed back into the refined 80S ribosomes using rRNA and ribosomal proteins as a guide, further real-space refined using Phenix v. dev-3885, and validated using Phenix validation tools which include the Molprobity suite (Williams et al., 2018). The deposited structures were obtained after real space refinement and model validation in Phenix v. dev-3885 (Afonine et al., 2018), as well as removing clashes > 1.0 in Coot v. 0.9-pre EL, part of the CCP-EM suite (Burnley et al., 2017). Figures were generated using Chimera v. 1.10.2 (Pettersen et al., 2004), Chimera X 1.0, and Pymol v. 1.8.0.5 and v. 2.0.6. (Schrödinger). Validation statistics are in Table S3.

## QUANTIFICATION AND STATISTICAL ANALYSIS

All *in vitro* experiments were repeated at least three times, and representative gel images and sucrose density gradient graphs were shown.

## DATA AND CODE AVAILABILITY

The atomic coordinates of the rotated and unrotated 80S/IRES complexes have been deposited in the Protein Data Bank (PDB). The accession numbers for the atomic models of the [80S-HalV IGR IRES] in the rotated and unrotated conformation reported in this paper are: 7A01 and 6ZVK, respectively.

The cryo-EM maps of the rotated and unrotated 80S/IRES complexes have been deposited in the Electron Microscopy Data Bank (EMDB) with the accession codes: EMD-11590 and EMD-11459, respectively. Gel images are available at: https://data.mendeley.com/datasets/wx2ypvtxh8/draft?a=8017d354-6b04-4735-a5f8-464205fbbad9

## REFERENCES

Abaeva, I.S., Pestova, T.V., and Hellen, C.U. (2016). Attachment of ribosomal complexes and retrograde scanning during initiation on the Halastavi árva virus IRES. Nucleic Acids Res. 44, 2362–2377.

Abeyrathne, P.D., Koh, C.S., Grant, T., Grigorieff, N., and Korostelev, A.A. (2016). Ensemble cryo-EM uncovers inchworm-like translocation of a viral IRES through the ribosome. Elife 5, pii: e14874.

Acosta-Reyes, F., Neupane, R., Frank, J., and Fernández, I.S. (2019). The Israeli acute paralysis virus IRES captures host ribosomes by mimicking a ribosomal state with hybrid tRNAs. EMBO J. 38, e102226.

Adams, P.D., Afonine, P.V., Bunkóczi, G., Chen, V.B., Davis, I.W., Echols, N., Headd, J.J., Hung, L.W., Kapral, G.J., Grosse-Kunstleve, R.W. et al. (2010). PHENIX: a comprehensive Python-based system for macromolecular structure solution. Acta Crystallogr. D Biol. Crystallogr. 66, 213–221.

Afonine, P., Headd, J., Terwilliger, T., and Adams, P. (2013). New tool: phenix. real_space_refine. Comp. Crystallog. Newsletter 4, 43–44.

Alkalaeva, E.Z., Pisarev, A.V., Frolova, L.Y., Kisselev, L.L., and Pestova, T.V. (2006). In vitro reconstitution of eukaryotic translation reveals cooperativity between release factors eRF1 and eRF3. Cell 125, 1125–1136.

Anger, A.M., Armache, J.P., Berninghausen, O., Habeck, M., Subklewe, M., Wilson, D.N., and Beckmann, R. (2013). Structures of the human and Drosophila 80S ribosome. Nature 497, 80–85.

Asnani, M., Kumar, P., and Hellen CU. (2015. Widespread distribution and structural diversity of type IV IRESs in members of Picornaviridae. Virology 478, 61–74.

Boros, Á., Pankovics, P., Simmonds, P., and Reuter, G. (2011). Novel positive-sense, single-stranded RNA (+ssRNA) virus with di-cistronic genome from intestinal content of freshwater carp (Cyprinus carpio). PLoS One 6, e29145.

Brown, A., Shao, S., Murray, J., Hegde, R.S., and Ramakrishnan, V. (2015). Structural basis for stop codon recognition in eukaryotes. Nature 524, 493–496.

Brown, A., Baird, M.R., Yip, M.C., Murray, J., and Shao, S. (2018). Structures of translationally inactive mammalian ribosomes. Elife 7, pii: e40486.

Budkevich, T.V., Giesebrecht, J., Behrmann, E., Loerke, J., Ramrath, D.J., Mielke, T., Ismer, J., Hildebrand, P.W., Tung, C.S., Nierhaus, K.H., Sanbonmatsu, K.Y., and Spahn, C.M. (2014). Regulation of the mammalian elongation cycle by subunit rolling: a eukaryotic-specific ribosome rearrangement. Cell 158, 121–131.

Chou, F.C., Sripakdeevong, P., Dibrov, S.M., Hermann, T., and Das, R. (2013). Correcting pervasive errors in RNA crystallography through enumerative structure prediction. Nat. Methods 10, 74–76.

Costantino, D., and Kieft, J.S. (2005). A preformed compact ribosome-binding domain in the cricket paralysis-like virus IRES RNAs. RNA 11, 332–343.

Costantino, D.A., Pfingsten, J.S., Rambo, R.P., and Kieft, J.S. (2008). tRNA-mRNA mimicry drives translation initiation from a viral IRES. Nat. Struct. Mol. Biol. 15, 57–64.

Culley, A.I., Lang, A.S., and Suttle, C.A. (2007). The complete genomes of three viruses assembled from shotgun libraries of marine RNA virus communities. Virology J. 4, 69.

de Breyne, S., Yu, Y., Unbehaun, A., and Pestova, T.V., and Hellen, C.U. (2009). Direct functional interaction of initiation factor eIF4G with type 1 internal ribosomal entry sites. Proc. Natl. Acad. Sci. USA 106, 9197–9202.

Dereeper, A., Guignon, V., Blanc, G., Audic, S., Buffet, S., Chevenet, F., Dufayard, J.F., Guindon, S., Lefort, V., Lescot, M. et al. (2008). Phylogeny.fr: robust phylogenetic analysis for the non-specialist. Nucleic Acids Res. 36, W465–469.

Emsley, P., and Cowtan, K. (2004). Coot: model-building tools for molecular graphics. Acta Crystallogr. D Biol. Crystallogr. 60, 2126–2132.

Fernández, I.S., Bai, X.C., Murshudov, G., Scheres, S.H., and Ramakrishnan, V. (2014). Initiation of translation by cricket paralysis virus IRES requires its translocation in the ribosome. Cell 157, 823–831.

Flis, J., Holm, M., Rundlet, E.J., Loerke, J., Hilal, T., Dabrowski, M., Bürger, J., Mielke, T., Blanchard, S.C., Spahn, C.M.T., and Budkevich, T.V. (2018). tRNA translocation by the eukaryotic 80S ribosome and the impact of GTP hydrolysis. Cell Rep. 25, 2676–2688.e7.

Heaton, J.H., Dlakic, W.M., Dlakic, M., and Gelehrter, T.D. (2001). Identification and cDNA cloning of a novel RNA-binding protein that interacts with the cyclic nucleotide-responsive sequence in the Type-1 plasminogen activator inhibitor mRNA. J. Biol. Chem. 276, 3341–3347.

Hoang, D.T., Chernomor, O., von Haeseler, A., Minh, B.Q., and Vinh, S.L (2018). UFBoot2: Improving the ultrafast bootstrap approximation. Mol. Biol. Evol. 35, 518–522.

Humphrey, W., Dalke, A., and Schulten, K. (1996). VMD: visual molecular dynamics. J. Mol. Graph. 14, 33–8, 27-8.

Imai, S., Kumar, P., Hellen, C. U., D’Souza, V. M., and Wagner, G. (2016). An accurately preorganized IRES RNA structure enables eIF4G capture for initiation of viral translation. Nat. Struct. Mol. Biol. 23, 859–864.

Jackson, R.J., Hellen, C.U., and Pestova, T.V. (2010). The mechanism of eukaryotic translation initiation and principles of its regulation. Nat. Rev. Mol. Cell. Biol. 11, 113–127.

Jain, S., Richardson, D.C., and Richardson, J.S. (2015). Computational methods for RNA structure validation and improvement. Methods Enzymol. 558, 181–212.

Jan, E. (2006). Divergent IRES elements in invertebrates. Virus Res. 119, 16–28.

Jan, E., and Sarnow, P. (2002). Factorless ribosome assembly on the internal ribosome entry site of cricket paralysis virus. J. Mol. Biol. 324, 889–902.

Jan, E., Kinzy, T.G., and Sarnow, P. (2003). Divergent tRNA-like element supports initiation, elongation, and termination of protein biosynthesis. Proc. Natl. Acad. Sci. USA 100, 15410–15415.

Jang, C.J., Lo, M.C., and Jan, E. (2009). Conserved element of the dicistrovirus IGR IRES that mimics an E-site tRNA/ribosome interaction mediates multiple functions. J. Mol. Biol. 387, 42–58.

Jang, C.J., and Jan, E. (2010). Modular domains of the Dicistroviridae intergenic internal ribosome entry site. RNA 16, 1182–1195.

Janssen, S., and Giegerich, R. (2015). The RNA shapes studio. Bioinformatics 31, 423–425.

Jossinet, F., Ludwig, T.E., and Westhof, E. (2010. Assemble: an interactive graphical tool to analyze and build RNA architectures at the 2D and 3D levels. Bioinformatics 26, 2057–2059.

Juneau, K., Podell, E., Harrington, D.J., and Cech, T.R. (2001). Structural basis of the enhanced stability of a mutant ribozyme domain and a detailed view of RNA--solvent interactions. Structure 9, 221–231

Kanamori, Y., and Nakashima, N. (2001). A tertiary structure model of the internal ribosome entry site (IRES) for methionine-independent initiation of translation. RNA 7, 266–274.

Kerr, C.H., Ma, Z.W., Jang, C.J., Thompson, S.R., and Jan, E. (2016). Molecular analysis of the factorless internal ribosome entry site in Cricket Paralysis virus infection. Sci. Rep. 6, 37319.

Koh, C.S., Brilot, A.F., Grigorieff, N., and Korostelev, A.A. (2014). Taura syndrome virus IRES initiates translation by binding its tRNA-mRNA-like structural element in the ribosomal decoding center. Proc. Natl. Acad. Sci. USA 111, 9139–9144.

Kucukelbir, A., Sigworth, F.J., and Tagare, H.D. (2014). Quantifying the local resolution of cryo-EM density maps. Nat. Methods. 11, 63–65.

Korostelev, A., Trakhanov, S., Laurberg, M., and Noller, H.F. (2006). Crystal structure of a 70S ribosome-tRNA complex reveals functional interactions and rearrangements. Cell 126, 1065–1077.

Lee, Y.J., Wei, H.M., Chen, L.Y., and Li, C. (2014). Localization of SERBP1 in stress granules and nucleoli. FEBS J. 281, 352–364.

Lomakin, I.B., Hellen, C.U., and Pestova, T.V. (2000). Physical association of eukaryotic initiation factor 4G (eIF4G) with eIF4A strongly enhances binding of eIF4G to the internal ribosomal entry site of encephalomyocarditis virus and is required for internal initiation of translation. Mol. Cell. Biol. 20, 6019–6029.

Lomakin, I.B., Shirokikh, N.E., Yusupov, M.M., Hellen, C.U., and Pestova, T.V. (2006). The fidelity of translation initiation: reciprocal activities of eIF1, IF3 and YciH. EMBO J. 25, 196–210.

Mailliot, J. and Martin, F. (2017). Viral internal ribosomal entry sites: four classes for one goal. WIREs RNA 9, e1458.

Mohr, I., and Sonenberg, N. (2012). Host translation at the nexus of infection and immunity. Cell Host Microbe 12, 470–483.

Muhs, M., Hilal, T., Mielke, T., Skabkin, M.A., Sanbonmatsu, K.Y., Pestova, T.V., and Spahn, C.M. (2015). Cryo-EM of ribosomal 80S complexes with termination factors reveals the translocated cricket paralysis virus IRES. Mol. Cell 57, 422–432.

Muhs, M., Yamamoto, H., Ismer, J., Takaku, H., Nashimoto, M., Uchiumi, T., Nakashima, N., Mielke, T., Hildebrand, P.W., Nierhaus, K.H., and Spahn, C.M. (2011). Structural basis for the binding of IRES RNAs to the head of the ribosomal 40S subunit. Nucleic Acids Res. 39, 5264–5275.

Murray, J., Savva, C.G., Shin, B.S., Dever, T.E., Ramakrishnan, V, and Fernández, I.S. (2016). Structural characterization of ribosome recruitment and translocation by type IV IRES. Elife 5, pii: e13567.

Nakashima, N., and Uchiumi, T. (2008). Functional analysis of structural motifs in dicistroviruses. Virus Res. 139, 137–147.

Neubauer, C., Gao, Y.G., Andersen, K.R., Dunham, C.M., Kelley, A.C., Hentschel, J., Gerdes, K., Ramakrishnan, V., and Brodersen, D.E. (2009). The structural basis for mRNA recognition and cleavage by the ribosome-dependent endonuclease RelE. Cell 139, 1084–1095.

Nguyen, L.-T., Schmidt, H.A., von Haeseler, A., and Minh, B.Q. (2015). IQ-TREE: A fast and effective stochastic algorithm for estimating maximum likelihood phylogenies. Mol. Biol. Evol. 32, 268–274.

Nishiyama, T., Yamamoto, H., Shibuya, N., Hatakeyama, Y., Hachimori, A., Uchiumi, T., and Nakashima, N. (2003). Structural elements in the internal ribosome entry site of *Plautia stali* intestine virus responsible for binding with ribosomes. Nucleic Acids Res. 31, 2434–2442.

Okonechnikov, K., Golosova, O., Fursov, M. and UGENE team. (2012). Unipro UGENE: a unified bioinformatics toolkit. Bioinformatics 28, 1166–1167.

Pedersen, K., Zavialov, A.V., Pavlov, M.Y., Elf, J., Gerdes, K., and Ehrenberg, M. (2003). The bacterial toxin RelE displays codon-specific cleavage of mRNAs in the ribosomal A site. Cell 112, 131–140.

Pestova, T.V., Hellen, C.U., and Shatsky, I.N. (1996). Canonical eukaryotic initiation factors determine initiation of translation by internal ribosomal entry. Mol. Cell. Biol. 16, 6859–6869.

Pestova, T.V., Borukhov, S.I., and Hellen, C.U. (1998a). Eukaryotic ribosomes require initiation factors 1 and 1A to locate initiation codons. Nature 394, 854–859.

Pestova, T.V., Shatsky, I.N., Fletcher, S.P., Jackson, R.J., and Hellen, C.U. (1998b). A prokaryotic-like mode of cytoplasmic eukaryotic ribosome binding to the initiation codon during internal translation initiation of hepatitis C and classical swine fever virus RNAs. Genes Dev. 112, 67–83.

Pestova, T.V., and Hellen, C.U. (2003). Translation elongation after assembly of ribosomes on the Cricket paralysis virus internal ribosomal entry site without initiation factors or initiator tRNA. Genes Dev. 17, 181–186.

Petrov, A., Grosely, R., Chen, J., O’Leary, S.E., and Puglisi, J.D. (2016). Multiple parallel pathways of translation initiation on the CrPV IRES. Mol. Cell 62, 92–103.

Petrov, A.S., Bernier, C.R., Hsiao, C., Norris, A.M., Kovacs, N.A., Waterbury, C.C., Stepanov, V.G., Harvey, S.C., Fox, G.E., Wartell, R.M., et al. (2014). Evolution of the ribosome at atomic resolution. Proc. Natl. Acad. Sci. USA 111, 10251–10256.

Pettersen, E.F., Goddard, T.D., Huang, C.C., Couch, G.S., Greenblatt, D.M., Meng, E.C., and Ferrin, T.E. (2004). UCSF Chimera--a visualization system for exploratory research and analysis. J. Comput. Chem. 25, 1605–1612.

Pfingsten, J.S., Castile, A.E., and Kieft, J.S. (2010). Mechanistic role of structurally dynamic regions in Dicistroviridae IGR IRESs. J. Mol. Biol. 395, 205–217.

Pfingsten, J.S., Costantino, D.A., and Kieft, J.S. (2006). Structural basis for ribosome recruitment and manipulation by a viral IRES RNA. Science 314, 1450–1454.

Pfingsten, J.S., Costantino, D.A., and Kieft, J.S. (2007). Conservation and diversity among the threedimensional folds of the Dicistroviridae intergenic region IRESes. J. Mol. Biol. 370, 856–869.

Phillips, J.C., Braun, R., Wang, W., Gumbart, J., Tajkhorshid, E., Villa, E., Chipot, C., Skeel, R.D., Kalé, L., and Schulten, K. (2005). Scalable molecular dynamics with NAMD. J. Comput. Chem. 26, 1781–1802.

Pisarev, A.V., Unbehaun, A., Hellen, C.U., and Pestova, T.V. (2007). Assembly and analysis of eukaryotic translation initiation complexes. Methods Enzymol. 430, 147–177.

Pisareva, V.P., Skabkin, M.A., Hellen, C.U., Pestova, T.V., and Pisarev, A.V. (2011). Dissociation by Pelota, Hbs1 and ABCE1 of mammalian vacant 80S ribosomes and stalled elongation complexes. EMBO J., 30, 1804–1817.

Pisareva, V.P., Pisarev, A.V., and Fernández, I.S. (2018). Dual tRNA mimicry in the Cricket paralysis virus IRES uncovers an unexpected similarity with the Hepatitis C virus IRES. Elife 7, pii: e34062.

Ruehle, M.D., Zhang, H., Sheridan, R.M., Mitra, S., Chen, Y., Gonzalez, R.L., Cooperman, B.S., and Kieft, J.S. (2015). A dynamic RNA loop in an IRES affects multiple steps of elongation factor-mediated translation initiation. Elife 4, pii: e08146.

Sasaki, J., and Nakashima, N. (2000). Methionine-independent initiation of translation in the capsid protein of an insect RNA virus. Proc. Natl. Acad. Sci. USA 97, 1512–1515.

Sato, K., Hamada, M., Asai, K., and Mituyama, T. (2009). CENTROIDFOLD: a web server for RNA secondary structure prediction. Nucleic Acids Res. 37, W277–W280.

Schüler, M., Connell, S.R., Lescoute, A., Giesebrecht, J., Dabrowski, M., Schroeer, B., Mielke, T., Penczek, P.A., Westhof, E., and Spahn, C.M. (2006). Structure of the ribosome-bound cricket paralysis virus IRES RNA. Nat. Struct. Mol. Biol. 13, 1092–1096.

Seit-Nebi, A., Frolova, L., Justesen, J., and Kisselev, L. (2001). Class-1 translation termination factors: invariant GGQ minidomain is essential for release activity and ribosome binding but not for stop codon recognition. Nucleic Acids Res. 29, 3982–3987.

Selmer, M., Dunham, C.M., Murphy, F.V. 4th, Weixlbaumer, A., Petry, S., Kelley, A.C., Weir, J.R., and Ramakrishnan, V. (2006). Structure of the 70S ribosome complexed with mRNA and tRNA. Science 313, 1935–1942.

Shi, M., Lin, X.D., Tian, J.H., Chen, L.J., Chen, X., Li, C.X., Qin, X.C., Li, J., Cao, J.P., Eden, J.S., et al. (2016). Redefining the invertebrate RNA virosphere. Nature 540, 539–543.

Skabkin, M.A., Skabkina, O.V., Hellen, C.U., and Pestova, T.V. (2013). Reinitiation and other unconventional posttermination events during eukaryotic translation. Mol. Cell 51, 249–264.

Trabuco, L.G., Villa, E., Mitra, K., Frank, J., and Schulten, K. (2008). Flexible fitting of atomic structures into electron microscopy maps using molecular dynamics. Structure 16, 673–683.

Wells, J.N., Buschauer, R., Mackens-Kiani, T., Best, K., Kratzat, H., Berninghausen, O., Becker, T., Gilbert, W., Cheng, J., and Beckmann, R. (2020). Structure and function of yeast Lso2 and human CCDC124 bound to hibernating ribosomes. PLoS Biol. 18, e3000780.

Williams, C.J., Headd, J.J., Moriarty, N.W., Prisant, M.G., Videau, L.L., Deis, L.N., Verma, V., Keedy, D.A., Hintze, B.J., Chen, V.B. et al. (2018). MolProbity: More and better reference data for improved allatom structure validation. Protein Sci. 27, 293–315.

Wilson, J.E., Pestova, T.V., Hellen, C.U., and Sarnow, P. (2000). Initiation of protein synthesis from the A site of the ribosome. Cell 102, 511–520.

Yamamoto, H., Nakashima, N., Ikeda, Y., and Uchiumi, T. (2007). Binding mode of the first aminoacyl-tRNA in translation initiation mediated by Plautia stali intestine virus internal ribosome entry site. J. Biol. Chem. 282, 7770–7776.

Zhu, J., Korostelev, A., Costantino, D.A., Donohue, J.P., Noller, H.F., and Kieft, J.S. (2011). Crystal structures of complexes containing domains from two viral internal ribosome entry site (IRES) RNAs bound to the 70S ribosome. Proc. Natl. Acad. Sci. USA 108, 1839–1844.

Zinoviev, A., Hellen, C.U., and Pestova, T.V. (2015). Multiple mechanisms of reinitiation on bicistronic calicivirus mRNAs. Mol. Cell 57, 1059–1073.

Zinoviev, A., Goyal, A., Jindal, S., LaCava, J., Komar, A.A., Rodnina, M.V., Hellen, C.U.T., and Pestova, T.V. (2018). Functions of unconventional mammalian translational GTPases GTPBP1 and GTPBP2. Genes Dev. 32, 1226–1241.

Zhang, K. (2016). Gctf: Real-time CTF determination and correction. J. Struct. Biol. 193, 1–12.

Zheng, S.Q., Palovcak, E., Armache, J.P., Verba, K.A., Cheng, Y., and Agard, D.A. (2017). MotionCor2: anisotropic correction of beam-induced motion for improved cryo-electron microscopy. Nat. Methods. 14, 331–332.

Zivanov, J., Nakane, T., Forsberg, B.O., Kimanius, D., Hagen, W.J., Lindahl, E., and Scheres, S.H. (2018). New tools for automated high-resolution cryo-EM structure determination in RELION-3. Elife 7, pii: e42166.

Zuker, M. (2003). Mfold web server for nucleic acid folding and hybridization prediction. Nucleic Acids Res. 31, 3406–3415.

