## Supplementary figures and tables for "The Halastavi árva virus intergenic region IRES promotes translation by the simplest possible initiation mechanism"

**SUPPLEMENTAL DATA**

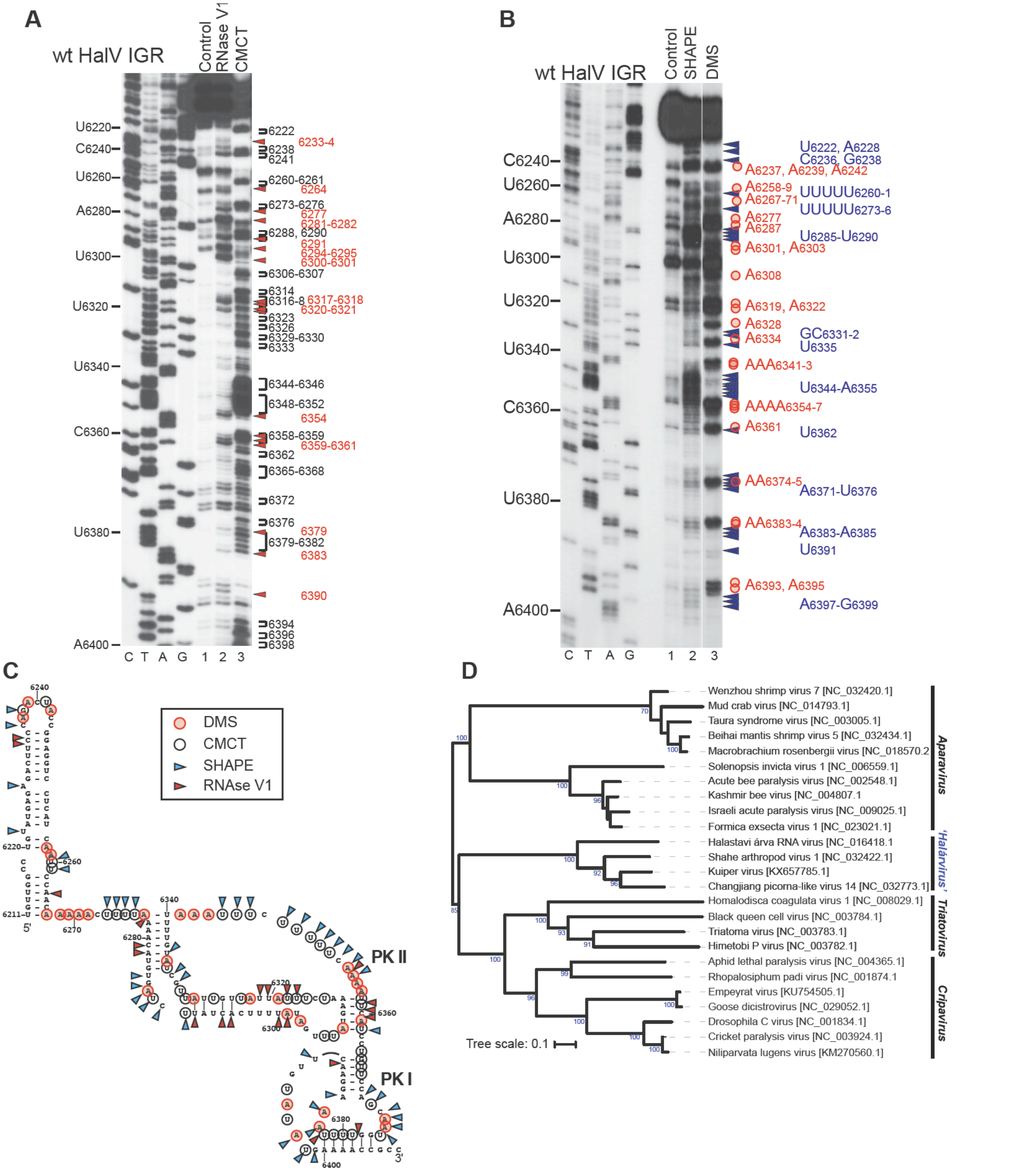


**Figure S1. Chemical and enzymatic probing of the HalV IGR IRES and phylogenetic analysis of dicistroviruses, related to Figure 3** . (A, B) Enzymatic (RNase V1) and chemical (CMCT, DMS, SHAPE) probing of the HalV IGR and the adjacent 3'-terminal region of ORF1. The positions of cleaved/modified nucleotides are indicated on the right using symbols (Rnase V1 - red arrowheads, CMCT - black brackets, DMS - red circles, NMIA - blue arrowheads). (B) Separation of lanes by white lines indicates that they were juxtaposed from the same gel. (C) Model of the HalV IGR and the adjacent 3'-terminal region of ORF1, derived as described in Materials and Methods, and indicating the positions of nucleotides cleaved by RNase V1 (red arrowheads), or modified by CMCT (black circles), DMS (red circles) or NMIA (blue arrowheads) based on data shown in (A, B). (D) Phylogenetic tree of dicistroviruses, based on analysis of the ORF2 capsid protein precursors from the indicated viruses, and showing that Halastavi árva virus belongs to a clade that is distinct from the *Cripavirus* genus, the *Triatovirus* genus and the two clades of insect- and crustacean-infecting viruses in the *Aparavirus* genus of *Dicistroviridae*. Sequences were aligned using CLUSTALW, and the phylogenetic tree was inferred in IQ-TREE using the maximum-likelihood method with 10,000 ultrafast bootstrap replicates. The numbers at the branch nodes represent the bootstrap confidence levels (above 70). Bar, 0.1 amino acid substitutions per site. The accession number of each viral sequence is indicated in parentheses.

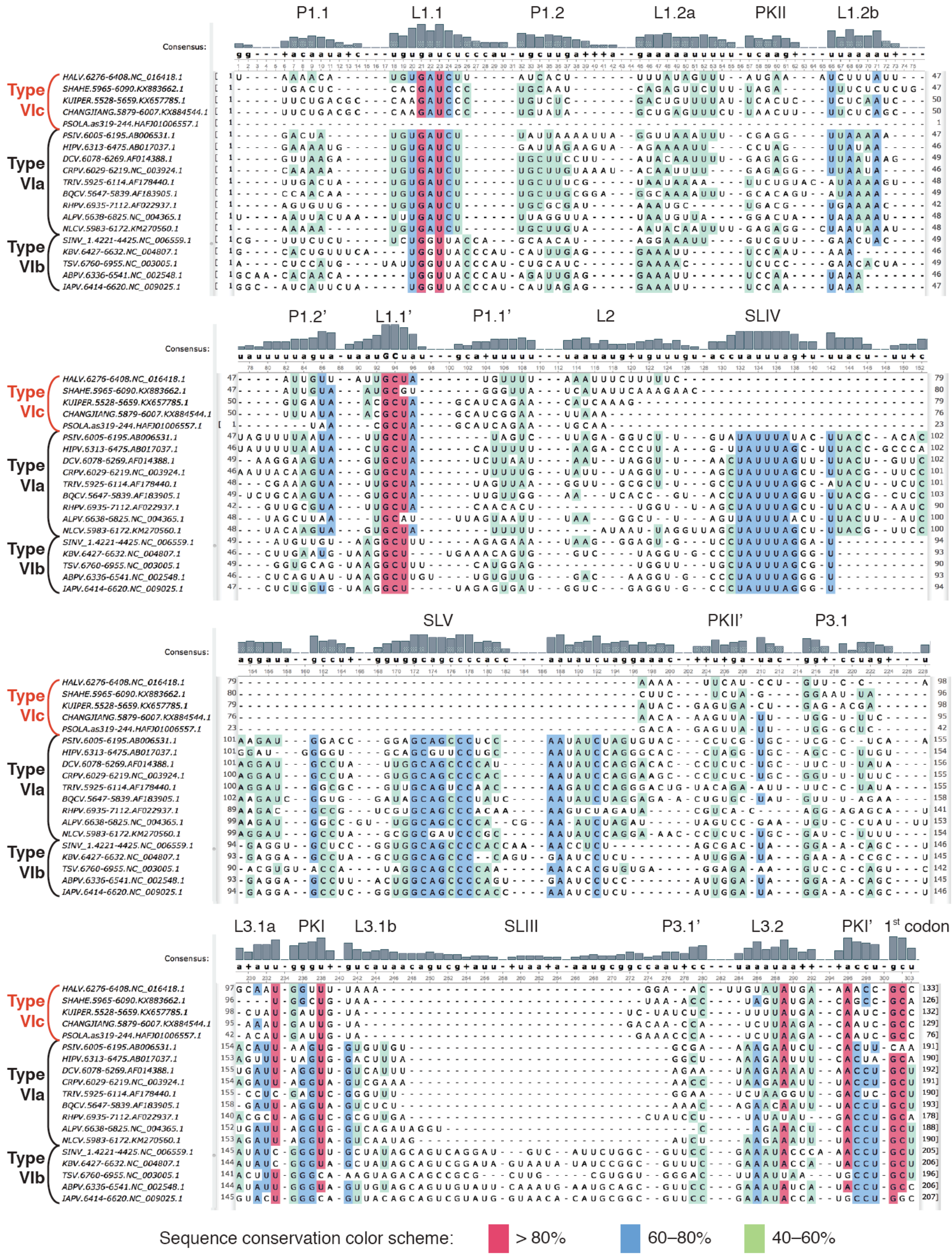


**Figure S2. Structure-based sequence alignment of representative type IV IGR IRES elements, related to Figures 3 and 4.**

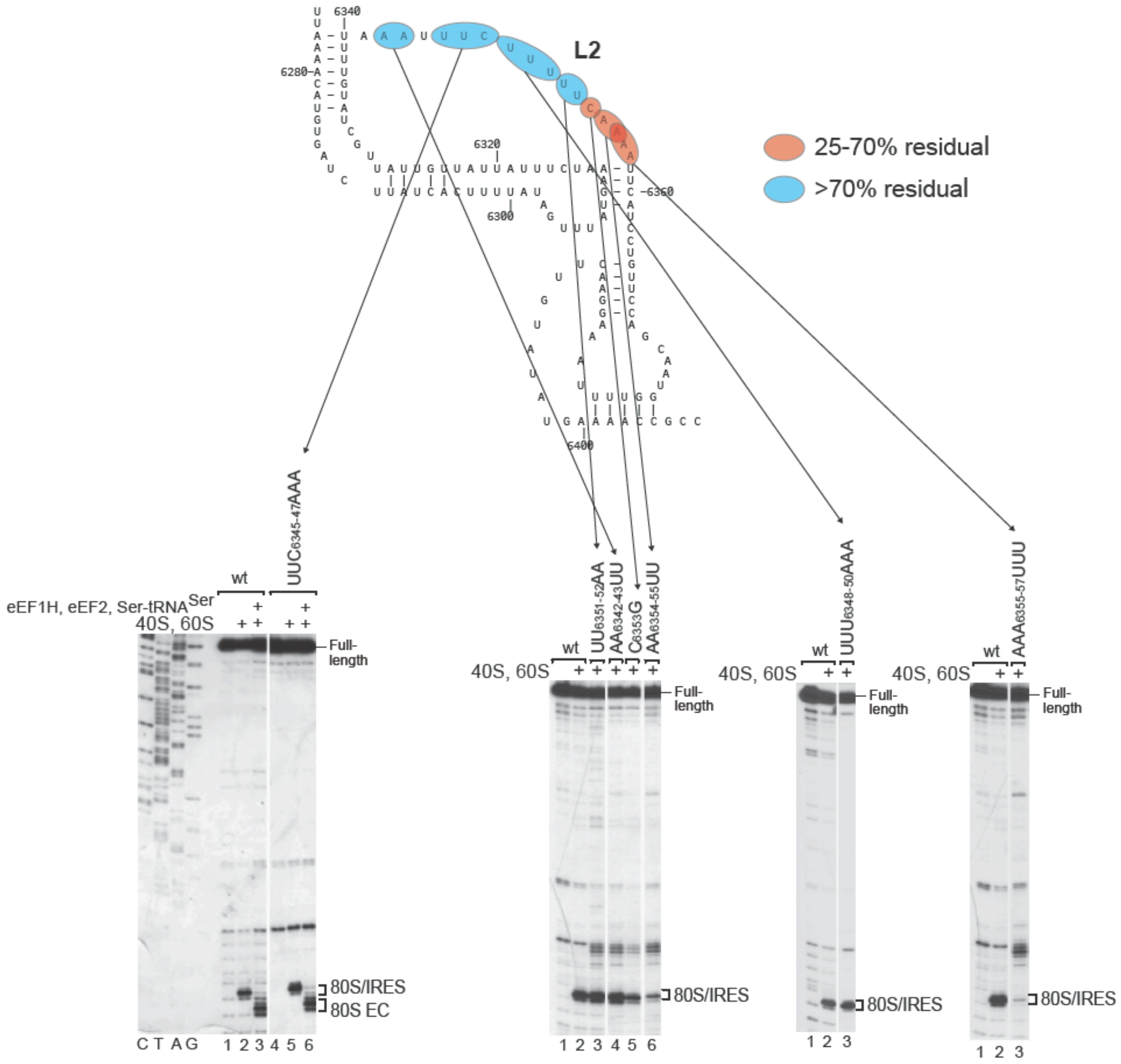


**Figure S3. Mutational analysis of the L2 loop of the HalV IGR IRES, related to Figure 3**. Activities of the HalV IGR L2 loop substitution mutants in ribosomal binding and one-cycle elongation, assayed by toe-printing and quantified by phosphorimager. Mutants were segregated into two classes based on their residual activity, mapped on the structural model of the IRES, and indicated by red shading (25-70% activity) and blue shading (>70% activity), respectively. Separation of lanes by white lines indicates that they were juxtaposed from the same gel.

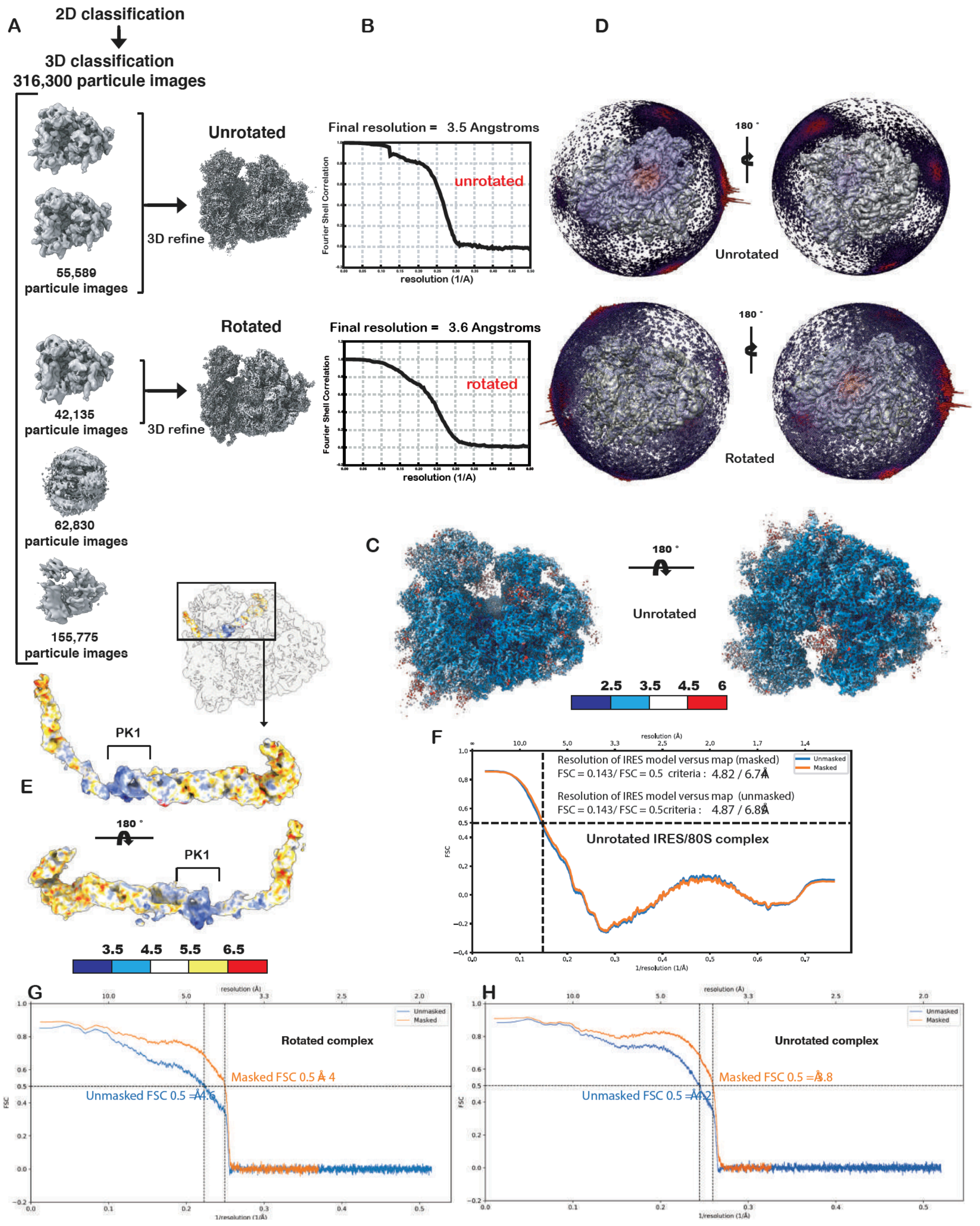


**Figure S4. Data processing workflow of 80S-HalV IRES data, local and model vs map resolution estimations, related to Figure 4.** (A) Graphical summary of the processing workflow described in Methods for the 80S-HalV complexes. Classification led to the identification of two rotational states of the 80S ribosomes, their average resolution FSC plots are presented in (B). (C) Local resolution of the unrotated 80S-HalV IRES complex. (D) Euler distribution plot for the unrotated and rotated reconstructions. (E) Local resolution map for the IRES region from the unrotated class reconstruction. (F) FSC curves of the HalV IRES atomic model vs its segmented map from the unrotated class reconstructions. (G) and (H) same as (F) but for the entire rotated and unrotated class reconstructions vs their atomic models. (F) to (H) are Phenix outputs from mtriage.

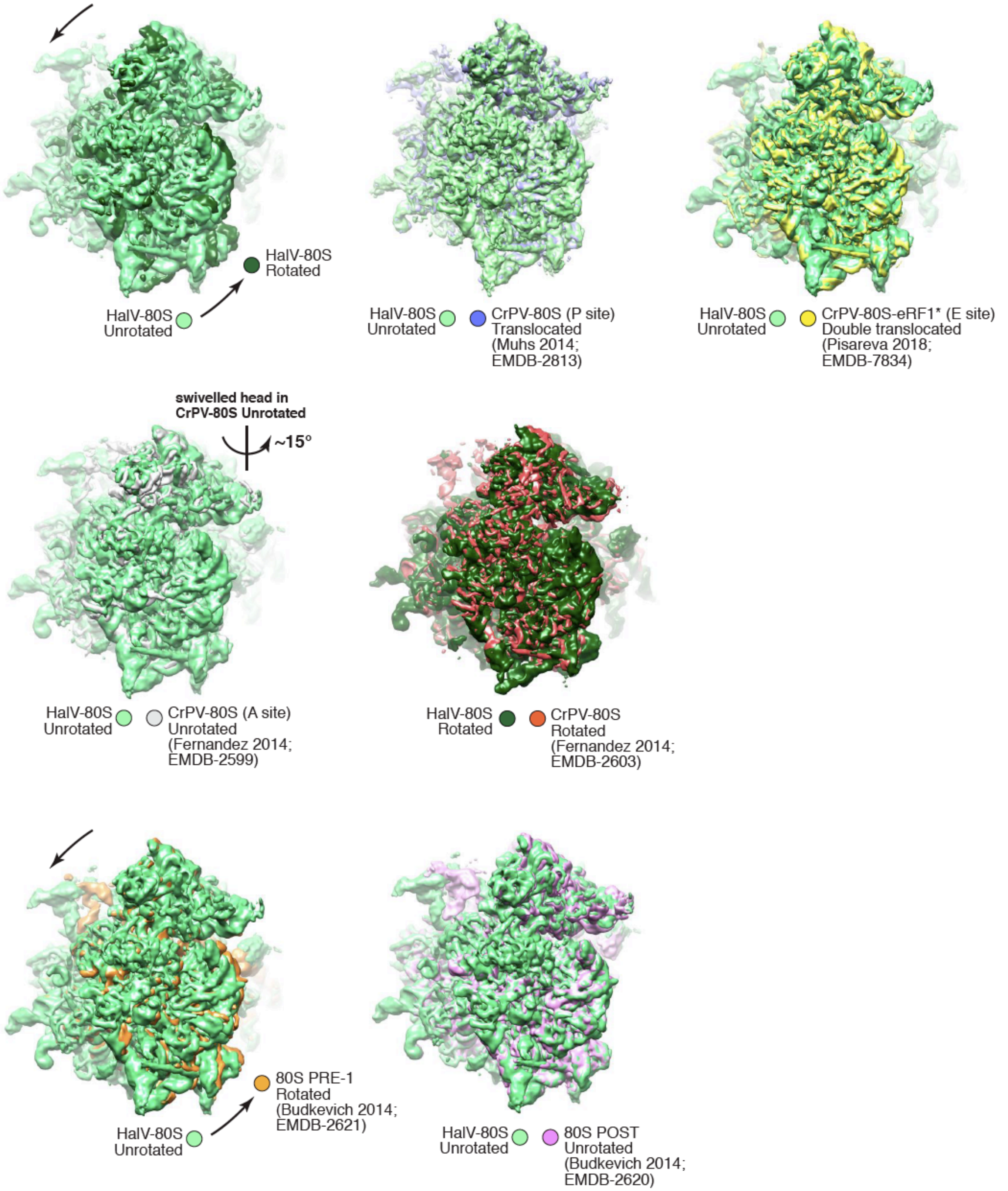


**Figure S5. Superimposition of experimental density maps for HalV-80S and other 80S-IGR IRES complexes, related to Figure 4**. Maps were Gaussian filtered to 1.5–2.5 for visual purposes. Superimpositions were based on the 60S subunit.

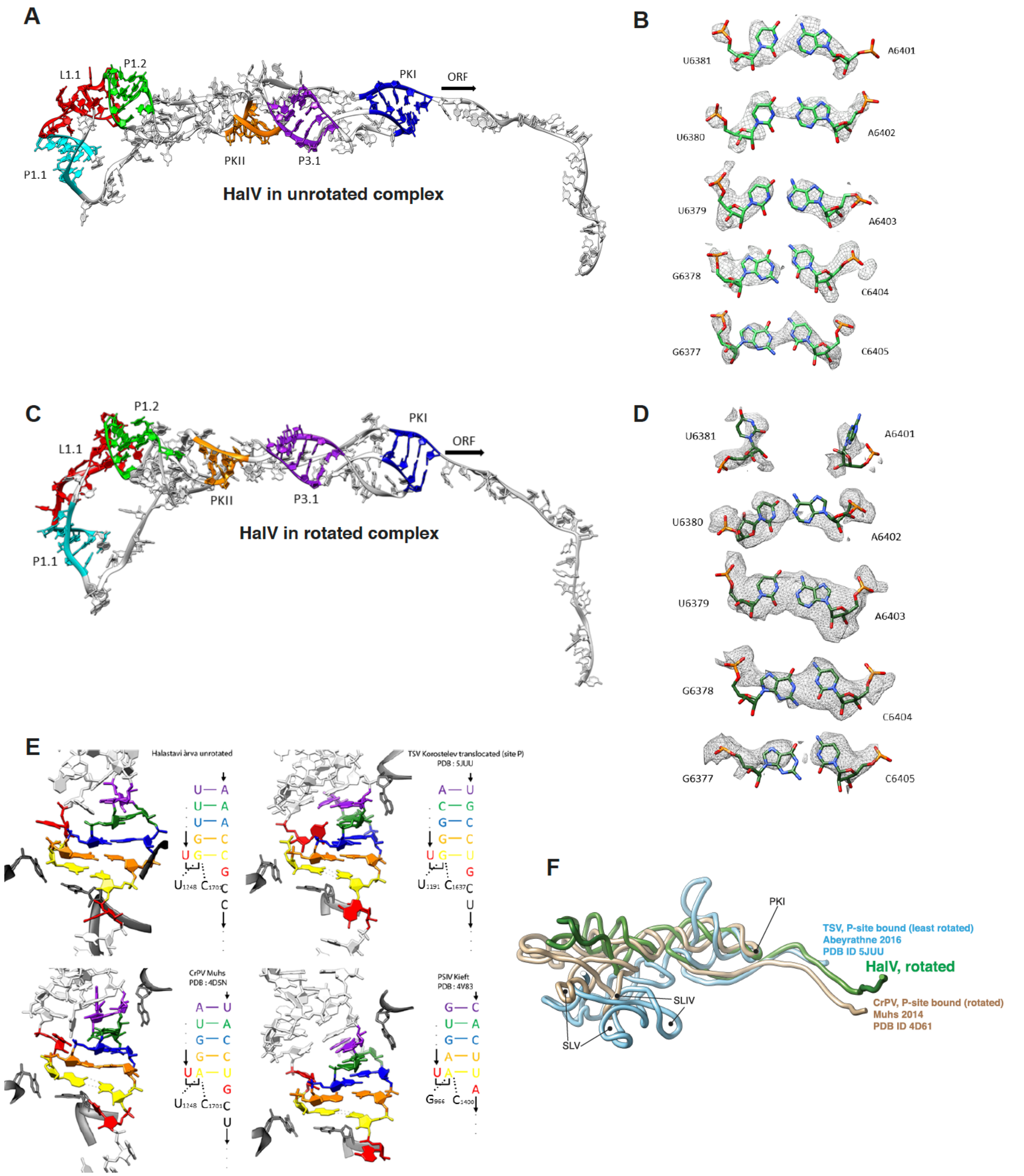


**Figure S6. Structural organization of the HalV IGR IRES and similarity with known structures, related to Figure 6.** (A) The IGR IRES in the unrotated state is color-coded according to helical elements. (B) Individual base pairs constituting PKI in the unrotated state (contour level 0.06). (C) Same as in (A) for the rotated state. (D) Same as (B) for the rotated state (contour level 0.08). (E) Color-coding of PKI per base pairs (including U...G in red that may be formed in some contexts). Shown are the unrotated HalV IGR IRES and other IGR IRESs in a similar post-translocation state. PKI typically contains five base pairs (colored in purple, green, blue, yellow, and orange). P-site 18S rRNA residues are shown in dark grey. (F) Superimposition of HalV and two other IGR IRESs, as indicated, based on the PKI region.

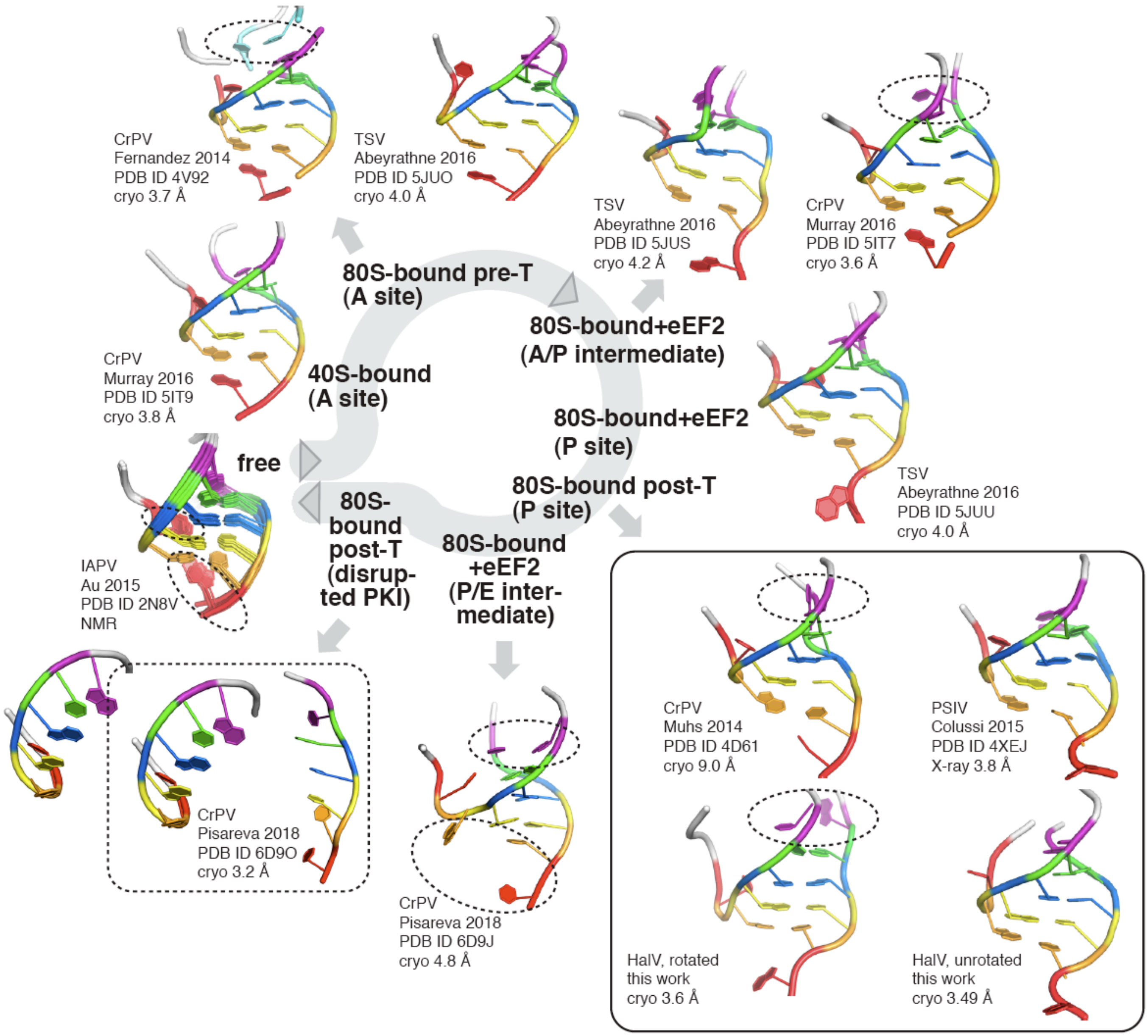


**Figure S7. Molecular details of PKI dynamics during the translation initiation/elongation cycle of IGR IRESs, related to Figure 6**. Same base-pair color-coding as in Figure S2. Base-pair disruption/formations are highlighted by dashed ovals. Post-translocation complexes with P-site bound IRESs are boxed. A disrupted PKI as revealed in a double-translocated complex (Pisareva et al., 2018) is in a dashed box. Legend to each structure: virus abbreviation, first author and year of publication, PDB ID, experimental method and resolution. Pre/post-translocation is abbreviated as pre/post-T.

**Supplemental Table 1. Amino acid sequence identity in the 3C protease/3D polymerase segment encoded by ORF1 of Halastavi árva virus (HalV) and related viruses.**

Percentage sequence identity determined by alignment using Clustal Omega of 3CD sequences from HalV, Shahe arthropod virus 1 (SAV1), Kuiper virus, Changjiang picorna-like virus 14 (CPLV14) and representative members of the genera *Cripavirus* (Cricket paralysis virus; CrPV), *Triatovirus* (Triatoma virus; TrV) and two clades of the genus *Aparavirus* (Taura syndrome virus (TSV) and Acute bee paralysis virus (ABPV) of *Dicistroviridae*. Sequence identity between 3CD moieties encoded by members of the proposed Halárvirus clade is indicated by bold text and yellow shading.

| | HalV | SAV1 | Kuiper | CPLV14 | TrV | TSV | CrPV | ABPV |
|---|---|---|---|---|---|---|---|---|
| HalV | **100.0** | - | - | - | - | - | - | - |
| SAV1 | **33.2** | **100.0** | - | - | - | - | - | - |
| Kuiper | **33.4** | **51.2** | **100.0** | - | - | - | - | - |
| CPLV14 | **32.7** | **52.6** | **71.2** | **100.0** | - | - | - | - |
| TrV | 23.7 | 24.2 | 23.2 | 23.8 | 100.0 | - | - | - |
| TSV | 26.9 | 25.0 | 25.4 | 25.0 | 25.2 | 100.0 | - | - |
| CrPV | 25.9 | 24.8 | 23.5 | 25.2 | 27.7 | 31.1 | 100.0 | - |
| ABPV | 27.4 | 25.9 | 27.3 | 26.9 | 29.4 | 33.3 | 34.5 | 100.0 |

**Supplemental Table 2. Amino acid sequence identity in the capsid protein precursor encoded by ORF2 of Halastavi árva virus (HalV) and related viruses.**

Percentage sequence identity determined by alignment using Clustal Omega of ORF2 sequences from HalV, Shahe arthropod virus 1 (SAV1), Kuiper virus, Changjiang picorna-like virus 14 (CPLV14) and representative members of the genera *Cripavirus* (Cricket paralysis virus; CrPV), *Triatovirus* (Triatoma virus; TrV) and two clades of the genus *Aparavirus* (Taura syndrome virus (TSV) and Acute bee paralysis virus (ABPV) of *Dicistroviridae*. Sequence identity between ORF2 moieties encoded by members of the proposed Halárvirus clade is indicated by bold text and yellow shading.

| | HalV | SAV1 | Kuiper | CPLV14 | TrV | TSV | CrPV | ABPV |
|---|---|---|---|---|---|---|---|---|
| HalV | **100.0** | - | - | - | - | - | - | - |
| SAV1 | **43.2** | **100.0** | - | - | - | - | - | - |
| Kuiper | **47.0** | **56.8** | **100.0** | - | - | - | - | - |
| CPLV14 | **44.4** | **53.2** | **60.2** | **100.0** | - | - | - | - |
| TrV | 22.0 | 22.4 | 22.6 | 22.0 | 100.0 | - | - | - |
| TSV | 20.8 | 21.3 | 22.2 | 22.3 | 22.3 | 100.0 | - | - |
| CrPV | 23.6 | 24.1 | 25.1 | 22.8 | 27.3 | 19.1 | 100.0 | - |
| ABPV | 21.2 | 24.5 | 22.0 | 23.4 | 23.7 | 23.2 | 21.5 | 100.0 |

**Supplemental table 3: Data collection and refinement statistics**

| Model | Unrotated | Rotated |
|---|---|---|
| ***Data collection and EM reconstruction*** | | |
| Microscope | FEI Titan Krios | |
| Voltage (kV) | 300 | |
| Camera | Falcon II direct electron detector | |
| Magnification (nominal) | 127,272 X | |
| Defocus range (*μ*m) | 0.6 – 4.5 | |
| Calibrated pixel size | 1.1 | |
| Electron exposure ($e^-$/Å$^2$) | 60 | |
| Exposure time (s) | 1.0 | |
| Number of frames per movie | 17 | |
| Automation software | EPU | |
| Number of micrographs | 2800 | |
| Initial particle number | 337268 | |
| Final particle number | 55589 | 42135 |
| Estimated accuracy of translation | 0.574 | 0.659 |
| Estimated accuracy of rotations | 0.524 | 0.718 |
| Map sharpening B factor (Å$^2$) | 154.9 | 153.8 |
| Map resolution (FSC = 0.143) | 3.49 | 3.6 |
| ***Refinement*** | | |
| **Composition (#)** | | |
| Chains | 86 | 86 |
| Atoms | 228292 (Hydrogen atoms: 0) | 224884 (Hydrogen atoms: 0) |
| Residues (Amino acids) | 11850 | 11858 |
| Residues (Nucleotides) | 6218 | 6055 |
| Water | 0 | 0 |
| Ligands (Type) | 6 (ZN) | 6 (ZN) |
| **Bonds (RMSD)** | | |
| Length (Å) (# > 4σ) | 0.006 (5) | 0.007 (9) |
| Angles (Å) (# > 4σ) | 0.845 (45) | 0.939 (186) |
| **MolProbity score** | 2.26 | 2.56 |
| **Clash score** | 15.55 | 25.85 |
| **EMRinger score** (scanned amino acids) | 2.06 (7532) | 0.94 (7534) |
| **Ramachandran plot (%)** | | |
| Outliers | 0.21 | 0.15 |
| Allowed | 10.54 | 14.54 |
| Favored | 89.26 | 85.31 |
| **Rama-Z (Ramachandran plot Z-score, RMSD)** | | |
| whole (N = 11690) | 3.42 (0.07) | 3.58 (0.07) |
| helix (N = 3584) | 2.24 (0.08) | 1.65 (0.08) |
| sheet (N = 1627) | 2.14 (0.12) | 1.86 (0.13) |
| loop (N = 6479) | 2.37 (0.07) | 3.04 (0.07) |
| **Rotamer outliers (%)** | 0.48 | 0.03 |
| **Cß outliers (%)** | 0.01 | 0.00 |
| **Peptide plane (%)** | | |
| Cis proline/general | 0.2/0.1 | 0.2/0.1 |
| Twisted proline/general | 0.8/0.3 | 0.8/0.3 |
| **CaBLAM outliers (%)** | 6.16 | 7.00 |
| **ADP (B-factors) (Å$^2$)** | | |
| Iso/Aniso (#) | 228292/0 | 224888/0 |
| min/max/mean | | |
| Protein | 15.89/160.37/64.69 | 20.14/179.89/76.11 |
| Nucleotide | 14.21/229.90/87.51 | 23.78/275.39/108.60 |
| Ligand | 43.94/127.55/87.92 | 46.56/174.83/109.42 |
| **Occupancy** (mean) | 1.0 | 1.0 |

| | | |
|---|---|---|
| **Box** | | |
| Lengths (Å) | 304.70. 297.00, 300.30 | 298.10, 288.20, 290.40 |
| Angles (°) | 90.00, 90.00, 90.00 | 90.00, 90.00, 90.00 |
| **Resolution Estimates (Å)** | | |
| d 99 (full) | 3.9 (Masked)<br>3.8 (Unmasked) | 4.0 (Masked)<br>4.0 (Unmasked) |
| d model | 3.8 (Masked)<br>3.8 (Unmasked) | 3.9 (Masked)<br>3.9 (Unmasked) |
| d FSC model (0/0.143/0.5) | 3.7/3.8/3.9 (Masked)<br>3.8/3.8/4.1 (Unmasked) | 3.9/3.9/4.0 (Masked)<br>3.9/3.9/4.5 (Unmasked) |
| **Map** min/max/mean | -0.15/0.25/0.00 | -0.07/0.12/0.00 |
| **Model vs. Data** | | |
| CC (mask) | 0.79 | 0.74 |
| CC (box) | 0.66 | 0.69 |
| CC (peaks) | 0.61 | 0.61 |
| CC (volume) | 0.77 | 0.73 |
| Mean CC for ligands | 0.81 | 0.70 |
